# Model-guided design of single-cargo, dual-attractor gene circuits

**DOI:** 10.64898/2026.10.19.760260

**Authors:** Emma L. Peterman, Nicholas Nolan, Paulina M. Naydenkov, Diya R. Godavarti, Domitilla Del Vecchio, Kate E. Galloway

## Abstract

Dual-attractor gene networks program distinct states, guiding native cellular decision-making processes such as differentiation. Synthetic dual-attractor circuits (DACs) emulate native gene networks, directing cells to two different states starting from a single, genetically uniform population. However, existing DAC designs for mammalian cells often require multiple genetic cargoes, limiting their application to control cell fate in therapeutically relevant contexts. Here, we leverage model-guided tuning of compact biochemical and biophysical layers of regulation to program two cellular states using a single-cargo DAC. Through activator selection and layering microRNA-mediated mutual repression, we identify and resolve an imbalance between modules that biases states, allowing us to achieve balanced adoption of both states without relying on copy-number variation. Consolidating this optimized DAC into a single cargo, we use gene syntax to minimize bias, generating two balanced states from a single cargo of less than 11 kilobases. Overall, this work establishes design rules for combined biochemical and biophysical tuning of DACs to generate balanced cell states from a single-cargo design, paving the way for programming divergent cell-state trajectories and dual-fate systems.

## Introduction

Attractor states are a fundamental feature of biological regulatory networks, guiding cells through development and differentiation [1–3]. The long-term stability of attractor states is determined by the underlying gene regulatory network (GRN). Transient attractors can guide cells through intermediate states along differentiation trajectories that continue to evolve [2, 4]. Stable attractors establish robust, persistent cell-fate decisions and reinforce cell identity [1]. In native systems, attractor networks are important for organizing cell-state transitions and establishing cell identity.

Dual-attractor gene networks direct genetically identical cells to two distinct states, allowing cells to adopt different downstream fates (Figure 1A) [5]. These decision-making processes are characterized by expression of two mutually inhibitory genes [6–8]. Through extracellular signal transduction or stochastic fluctuations in expression levels, cells can adopt two different cell states, where expression of one gene dominates over the other. Each of these states corresponds to a distinct attractor on the gene expression landscape (Figure 1A) [9, 10]. These regulatory networks provide a conceptual basis for engineering populations in which cells reliably adopt alternative functional states.

**Figure 1.**
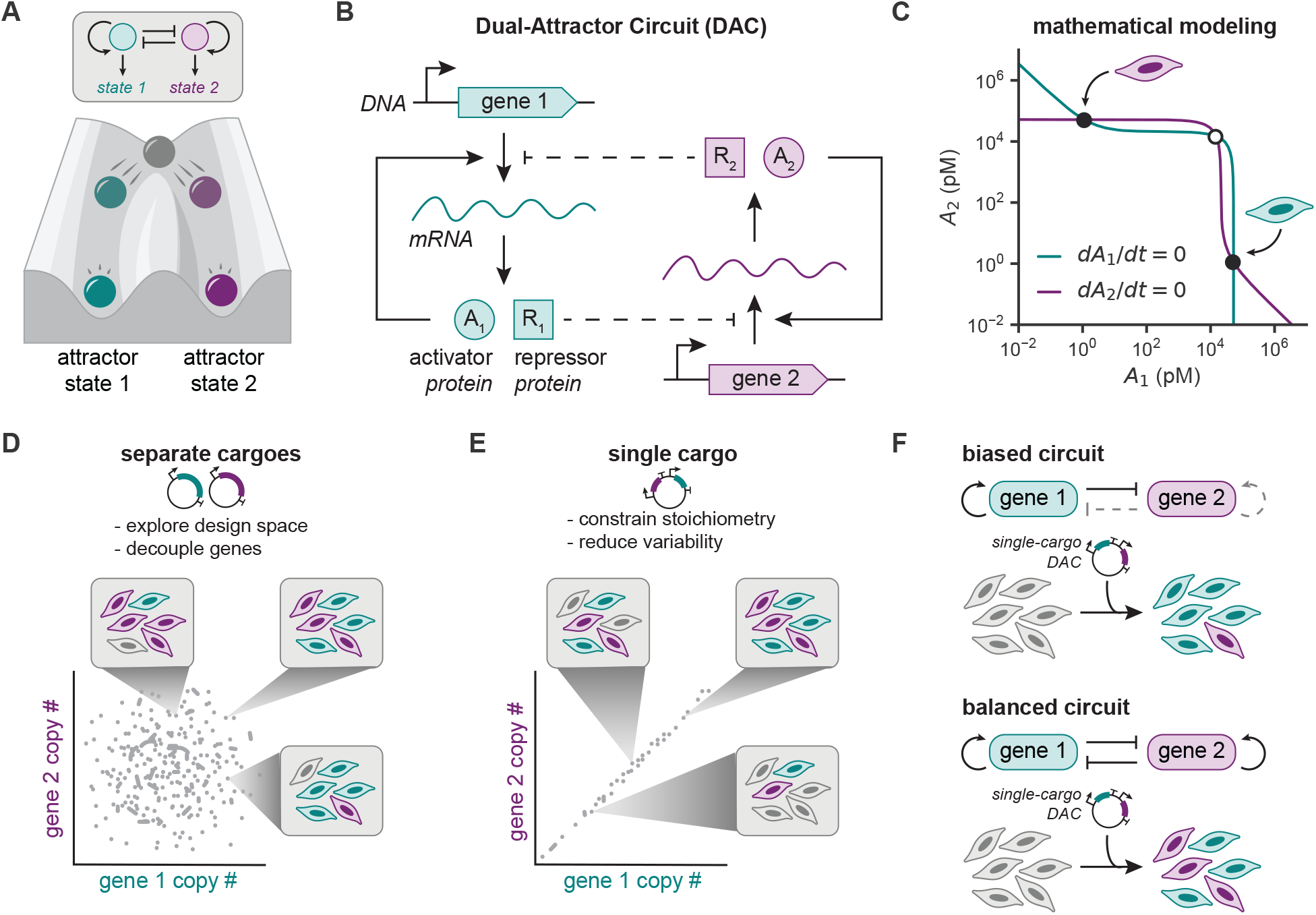
Dual attractor circuits (DACs) can program divergent cell states, but their translation is limited by delivery as multiple cargoes. (**A**) During development, bipotent progenitor cells can stochastically adopt one of two cell states, where each state represents a distinct attractor on the gene expression landscape. (**B**) In a gene circuit, the attractor states are programmed by two genes, each encoding a transcriptional activator (*A*_1_ or *A*_2_) and a transcriptional repressor (*R*_1_ or *R*_2_). Each activator can bind to its own promoter, implementing positive feedback. Each repressor can bind to the promoter of the opposite gene, inhibiting transcriptional activation and implementing mutual repression (dashed lines). (**C**) Nullcline analysis for the gene circuit depicted in **B** using the base parameters reported in Table S1. Filled circles indicate stable fixed points, and the unfilled circle indicates an unstable fixed point. The deterministic model is defined by Equations S1 to S10 as derived in the supplementary text. (**D**) Delivery of circuit components as separate cargoes enables rapid exploration of the circuit design space, making it a valuable research tool. However, differences in relative copy number can cause undesirable variability for downstream applications. (**E**) Delivery of the DAC as a single cargo constrains the stoichiometry of the genes, ensuring that all cells receive equal copy numbers of the genes. Nevertheless, this approach introduces biophysical coupling due to co-localization of the genes, adding an additional design variable to be optimized. (**F**) Existing dual-attractor circuit designs often exhibit a bias towards one of the two states. This results in significantly imbalanced cell-state commitment, making it difficult to engineer mixed populations with these circuits.

Synthetic biology has recreated dual-attractor behavior through the design of toggle switches [11–13] and other bistable circuits [14], demonstrating that two mutually exclusive states can be programmed synthetically. These dual-attractor circuits (DACs) program two distinct states that are defined by two exogenous mutually inhibitory genes [15, 16]. In this work, we selected a DAC architecture originally developed by Lebar *et al* [12]. This design relies on competitive binding between synthetic activator and repressor transcription factors (TFs) and enables the use of designable DNA-binding domains (DBDs) (Figure 1B). In the circuit, each transcriptional activator is expressed from a promoter containing its cognate target sites, implementing positive feedback. Additionally, each gene expresses a transcriptional repressor that binds to the promoter of the opposite gene, preventing binding of the transcriptional activator. Modeling this DAC using ordinary differential equations, we find two mathematically stable attractor states corresponding to each of the genes (Figure 1C). These align with experimentally measurable cell populations [12].

Synthetic DACs are often assembled from multiple genetic cargoes, producing variation in component copy number, integration site, and stoichiometry [17]. During circuit development, this variation can be leveraged for optimization. For example, techniques like poly-transfection allow for sampling of different component ratios [18]. This supports rapid exploration of the circuit design space and identification of stoichiometries that produce functional dual-attractor behavior (Figure 1D). However, this variation in relative copy number causes a reliance on monoclonal selection for homogenous performance [13]. Therefore, identifying rare functional clones limits the use of DACs in applications requiring reliable performance across heterogeneous, polyclonal cell populations, such as cell therapy engineering.

Single-cargo design of DACs has the potential to improve performance in polyclonal cell populations by enabling more homogeneous delivery of circuit components [17]. Once functional architectures and stoichiometries are identified, encoding all circuit components on a single cargo constrains their relative dosage and reduces variation arising from independent integration (Figure 1E). Therefore, single-cargo architectures offer a route toward reproducible dual-attractor behavior within polyclonal populations.

Nevertheless, implementation as a single cargo creates a new engineering problem. First, consolidating circuit components eliminates the ability to independently tune their copy numbers. Existing DAC designs can exhibit a bias towards one state (Figure 1F) [12–14], and these circuits are most commonly optimized by varying the relative copy number of each gene in the circuit and selecting a rare population of cells that exhibits the desired behavior [13]. Next, physical co-localization of genes can also introduce biophysical coupling and interference between regulatory units [19–21]. This means that designs optimized as separate cargoes may not retain their behavior when combined. Finally, single-cargo delivery imposes a constraint on total cargo size. For downstream compatibility with commonly used integration methods (e.g. lentivirus or PiggyBac transposase), the total DNA cargo size must be minimized, ideally under 10 kilobases (kb) for greatest efficiency [22–24]. Thus, successful single-cargo implementation requires understanding how architecture, stoichiometry, and regulatory coupling determine dual-attractor behavior.

The design rules required to build dual-attractor circuits on a single genetic cargo remain poorly defined. Establishing these rules is necessary to move dual-attractor circuits toward predictable, scalable function across polyclonal cell populations. In this work, we examine the molecular determinants of cell-state bias in a compact mammalian DAC based on transcriptional activation and repression. We identified imbalanced transcriptional activator strength as a key driver of circuit bias. To reduce the bias of the circuit without substantially increasing the circuit size or varying the relative copy number of each gene, we investigated two orthogonal and complementary methods of tuning the circuit biochemically: (1) varying the transcriptional activation domain and (2) layering additional microRNA-based mutual repression. Finally, we harnessed biophysical optimization of DACs by using gene syntax to reduce bias on a single cargo without expanding cargo size. Together, this work integrates experimental characterization and mechanistic modeling to reveal design rules for DAC composition and establishes a compact, single-cargo DAC design for mammalian cells.

## Results

### Mutual inhibition and positive feedback by ZF-based transcription factors generates two cell states

While the DAC design originally developed by Lebar *et al* established two states in HEK293T transfection [12], this circuit required delivery of four separate plasmids with a total cargo size of over 15 kb. This large cargo size is a barrier for translation of the circuit to other cell types and delivery methods. To significantly reduce the cargo size while using designable DBDs, we exchanged the large TALE domains in the original design for smaller zinc finger (ZF) domains. ZF domains have similar modularity and tunability to TALE domains but significantly shorter DNA sequences. Following the approach of Donahue *et al* [25], ZF-based transcriptional repression can be implemented by fusing a ZF domain to a fluorescent protein. These ZF-based repressors compete with ZF-based activators for binding sites and can prevent binding of adjacent activators through steric hindrance (Figure S2A). In HEK293T cell transfection, we find that ZF-based repression is dose-dependent, with reporter expression decreasing monotonically as a function of repressor expression level (Figure S2B). Importantly, repressors containing two different ZF domains, ZF37 and ZF43 [25], are orthogonal and only reduce reporter expression when the activator and repressor share matching ZF domains (Figure S2B). Since these repressors have distinct fluorescent markers and only one repressor is expected to be expressed in each state, the circuit state can be tracked via flow cytometry or imaging without needing to encode a separate fluorescent reporter (Figure 2A). Additionally, the smaller size of these TF sequences allows them to be encoded polycistronically using a 2A self-cleaving peptide. Together, this proposed DAC design can be programmed through delivery of two genes with a total sequence under 5 kb, representing an over 66% reduction in DNA cargo size compared to the original implementation (Figure 2A).

**Figure 2.**
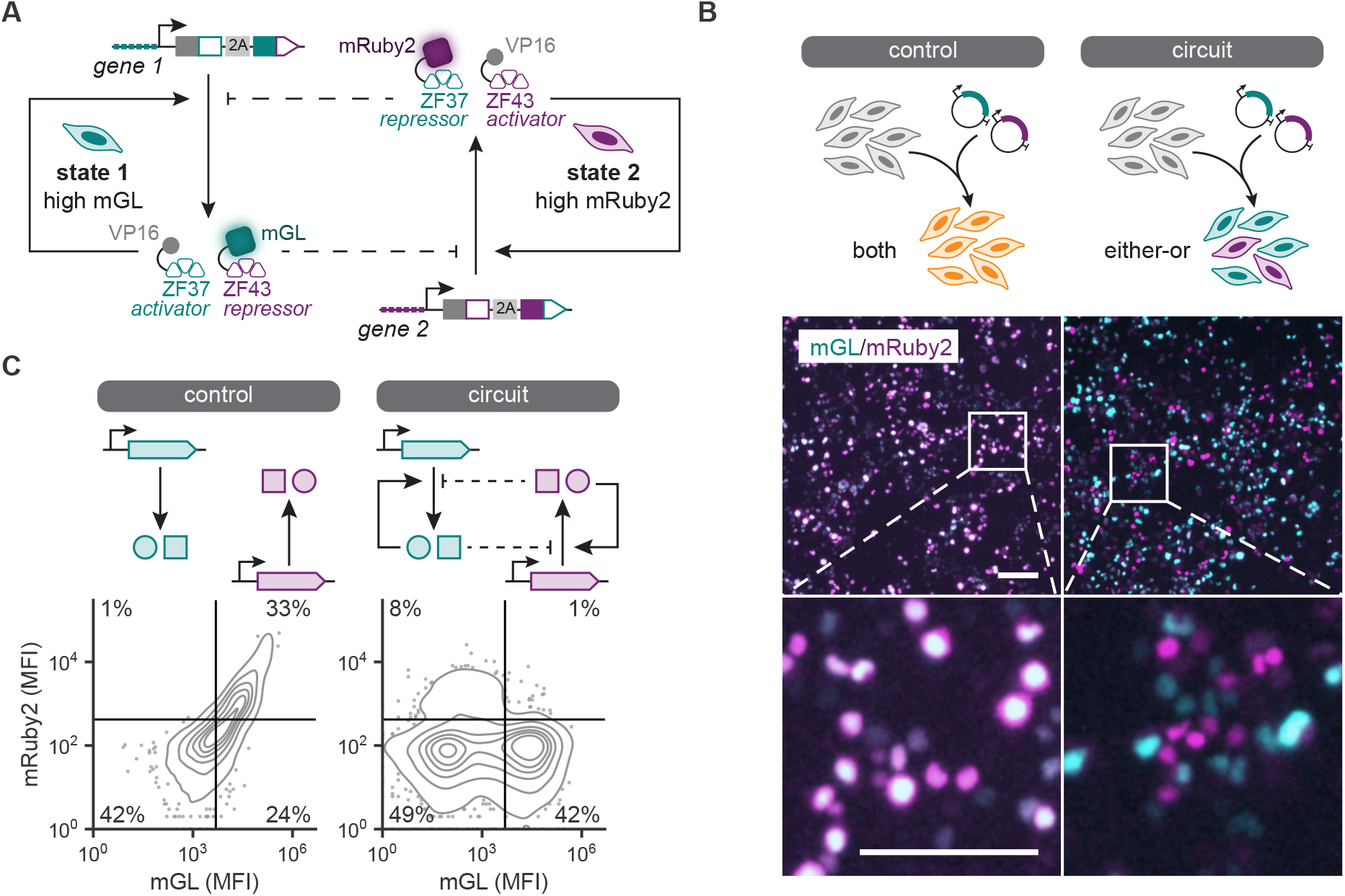
Mutual inhibition and positive feedback by ZF-based transcription factors generates two cell states. (A) Schematic of the ZF-based competitive binding circuit. Each gene is expressed via a ZF-responsive promoter containing 6 ZF binding sites upstream of the YB-TATA minimal promoter. Each gene encodes the transcriptional activator for its own promoter and the transcriptional repressor for the opposite promoter. State 1 is characterized by high expression of mGL (gene 1) and low expression of mRuby2 (gene 2). State 2 is characterized by high expression of mRuby2 (gene 2) and low expression of mGL (gene 1). The two circuit genes were co-transfected in HEK293T cells along with a constitutively expressed iRFP670 marker. The two genes in the control condition are each expressed from a constitutive EF1α promoter rather than a ZF-responsive promoter. Cells were analyzed by flow cytometry and imaging at 2 days post transfection (dpt). (**B**) Images of HEK293T cells co-transfected with two plasmids encoding either the control genes or circuit genes. Scale bars represent 100 µm. (**C**) Density plots of cells positive for the iRFP670 transfection marker for one representative biological replicate. Points represent the 5% of probability mass lying outside the lowest iso-proportion contour. All data is in arbitrary units from a flow cytometer. Gates are set based on poly-transfection characterization of these plasmid constructs using expression distributions in the lowest transfection marker bins. These data along with the corresponding gates are reported in Figures S4 and S5.

As an initial proof-of-concept, we tested the circuit in transient co-transfection of HEK293T cells, comparable to the original approach used by Lebar *et al* [12]. In this initial test, we delivered separate plasmids encoding the circuit genes as well as a fluorescent transfection marker in equal ratios. These co-transfected plasmids are delivered in roughly equal copy numbers to each cell, but the total copy number of plasmids taken up by cells can vary widely across the population [18]. In this context, expression of either circuit gene is driven initially by leaky expression from the ZF-responsive promoters. In transfection of HEK293T cells, this leakiness is sufficient for activation of the positive feedback loops due to the high copy number of the plasmids.

In the absence of positive feedback and mutual inhibition, the transfected cells exhibit correlated expression of both genes, which is consistent with co-transfection of reporter plasmids with no additional biochemical regulation (Figures 2B and 2C). In comparison, the DAC plasmids generate multiple populations (Figures 2B and 2C): (1) a negative population without detectable expression of either circuit gene, (2) an mGreenLantern-positive (mGL+) population corresponding to state 1, and (3) an mRuby2-positive (mRuby2+) population corresponding to state 2. Rather than expressing both circuit genes like in the control condition, expression of one circuit gene dominates, likely determined by the leaky activation kinetics of the two ZF-responsive promoters. We observe circuit activation in approximately 50% of transfected cells (Figure 2C), which aligns with previously published results for a circuit with minimal promoters [12]. Cells that do not activate either circuit gene despite expressing the transfection marker may have insufficient leaky expression due to low copy number. Additionally, the circuit displays a 5:1 bias in cells adopting state 1 over state 2 (Figure 2C), which is comparable to previously published results [12]. These results confirm that a DAC using ZF-mediated competitive binding performs comparably to the original TALE-based implementation while significantly reducing the circuit cargo size. However, more optimization is needed to reduce the bias of the circuit towards state 1 and identify methods to predictably tune the ratio of the two states.

### Varying relative copy number quantifies cell-state bias in a DAC

Copy number variation provides a window to quantitatively survey the bias of the circuit. To efficiently assess the circuit bias over a wide range of copy numbers, we performed poly-transfection experiments in which each circuit gene is prepared in separate transfection mixes with a second, constitutive marker plasmid (Figure 3A). Each constitutive marker (tagBFP or iRFP670) serves as a proxy for copy number for one of the circuit genes (gene 1 or gene 2, respectively). By preparing separate transfection mixes, the resulting copy numbers of transfected plasmids are not correlated between the two mixes, allowing for rapid sampling of a large copy number space [18]. Based on expression of the two constitutive markers, we separated cells into 25 copy number bins (Figures S3 and 3A). Within each bin, we can observe the distribution of mGL and mRuby2 expression (Figure S4) as well as calculate a bias score, defined as the difference between the numbers of mGL+ and mRuby2+ cells divided by their sum (Figure 3A). Perfect balance between the two states is represented by a bias score of zero. Positive scores represent a bias towards state 1 (mGL+), and negative scores represent a bias towards state 2 (mRuby2+). As a control, we confirmed that unimodal distributions are observed across all bins for a condition without positive feedback or mutual inhibition (Figure S5).

**Figure 3.**
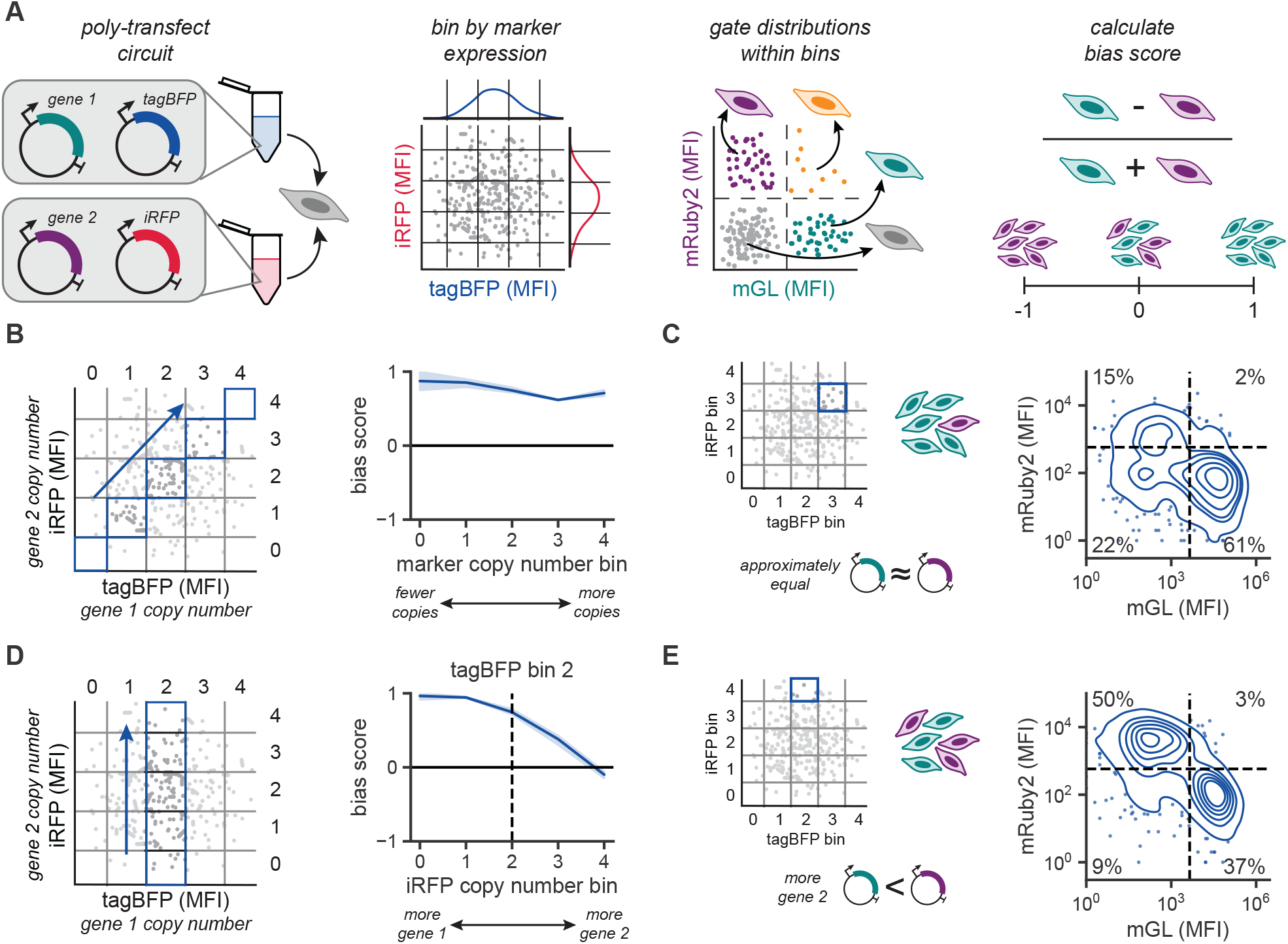
Varying relative copy number quantifies cell-state bias in a DAC. (**A**) Poly-transfection is used to survey a wide range of gene 1:gene 2 copy numbers [18]. Each co-transfection mix contains a marker plasmid constitutively expressing a fluorescent protein (tagBFP or iRFP670). At 2 dpt, cells are analyzed by flow cytometry and separated into equal width bins by tagBFP and iRFP670 expression. Within each bin, cells are gated into four quadrants based on mGL (gene 1) and mRuby2 (gene 2) expression. Finally, the bias score is calculated as the difference between the numbers of mGL+ and mRuby2+ cells divided by their sum. (**B**) Plot of gene circuit bias for the set of diagonal bins indicated in the left schematic. Data points represent the mean of three biological replicates, and shaded regions represent the 95% confidence interval. (**C**) Density plot of cells in tagBFP bin 3 and iRFP670 bin 3 for one representative biological replicate. Points represent the 5% of probability mass lying outside the lowest iso-proportion contour. Data is in arbitrary units from a flow cytometer. (**D**) Plot of gene circuit bias for varying iRFP670 bin while holding the tagBFP bin constant. Data points represent the mean of three biological replicates, and shaded regions represent the 95% confidence interval. Dashed black line corresponds to equal copy numbers of gene 1 and gene 2. (**E**) Density plot of cells in tagBFP bin 2 and iRFP670 bin 4 for one representative biological replicate. Points represent the 5% of probability mass lying outside the lowest iso-proportion contour. Data is in arbitrary units from a flow cytometer.

We first consider bins along the *y* = *x* diagonal, where the relative copy numbers of genes 1 and 2 are approximately equal but the total copy number varies (Figure 3B). These bins are most representative of co-transfection at equal ratios (Figure 2), where we previously observed a bias towards state 1. Similarly, the circuit exhibits a positive bias score across these diagonal bins, indicating a strong bias towards state 1 (Figure 3B). Looking at the distribution of mGL and mRuby2 expression within one of these diagonal bins, we again observe more cells in the mGL+ state when the two genes are delivered at approximately equal copy numbers (Figure 3C).

To probe the effects of varying relative copy number, we selected an intermediate gene 1 copy number bin (bin 2) and considered how the circuit bias changes with varying gene 2 copy number bin (Figure 3D). Indeed, we find that increasing the copy number of gene 2 relative to gene 1 reduces the bias of the circuit, and the least biased distribution is found with the highest gene 2 copy number (Figure 3D). Looking at the distribution of mGL and mRuby2 expression within this bin, we find much more balanced activation of both circuit states (Figure 3E). Therefore, state 2 (mRuby2+) is most robustly activated when gene 2 is present at a higher copy number than gene 1. While varying relative copy number is a simple, effective method of tuning cell-state ratios in transfection, this is not compatible with a single-cargo DAC design. To move towards translation of this technology, we instead explored two methods of optimizing cell-state ratios through biochemical tuning.

### Circuit bias can be reduced by balancing transcriptional activation strength

To understand the origin of our DAC bias, we characterized each transcriptional activator independently (Figure S6). Despite both original ZF activators having the same activation domain (VP16), these TFs induce different levels of reporter expression (Figure S6). We hypothesized that this imbalance in reporter activation could contribute to the bias observed when these TFs are used in the DAC. To address this, we characterized a panel of four additional activation domains of varying strengths (p65, VP64, VPR, and NFZ) (Figures 4A and S6). Using poly-transfection, we quantified the activation of a fluorescent reporter for varying levels of ZF activator expression (Figure S6). We then fit this experimental data to a Hill function to obtain a parameter, *α*, corresponding to the maximum reporter activation level (Figure 4A). While the VP16, p65, and VP64 domains all have significantly higher reporter activation when paired with ZF37, two activation domains—NFZ and VPR—induce strong and balanced activation when paired with either ZF37 or ZF43 (Figure 4A). This may indicate that the NFZ and VPR domains saturate HEK293T gene expression, obscuring any differences in activation due to ZF binding affinity. Therefore, we hypothesized that exchanging the VP16 activation domains in the circuit for NFZ or VPR domains could achieve more balanced activation of states 1 and 2 at one-to-one copy numbers.

**Figure 4.**
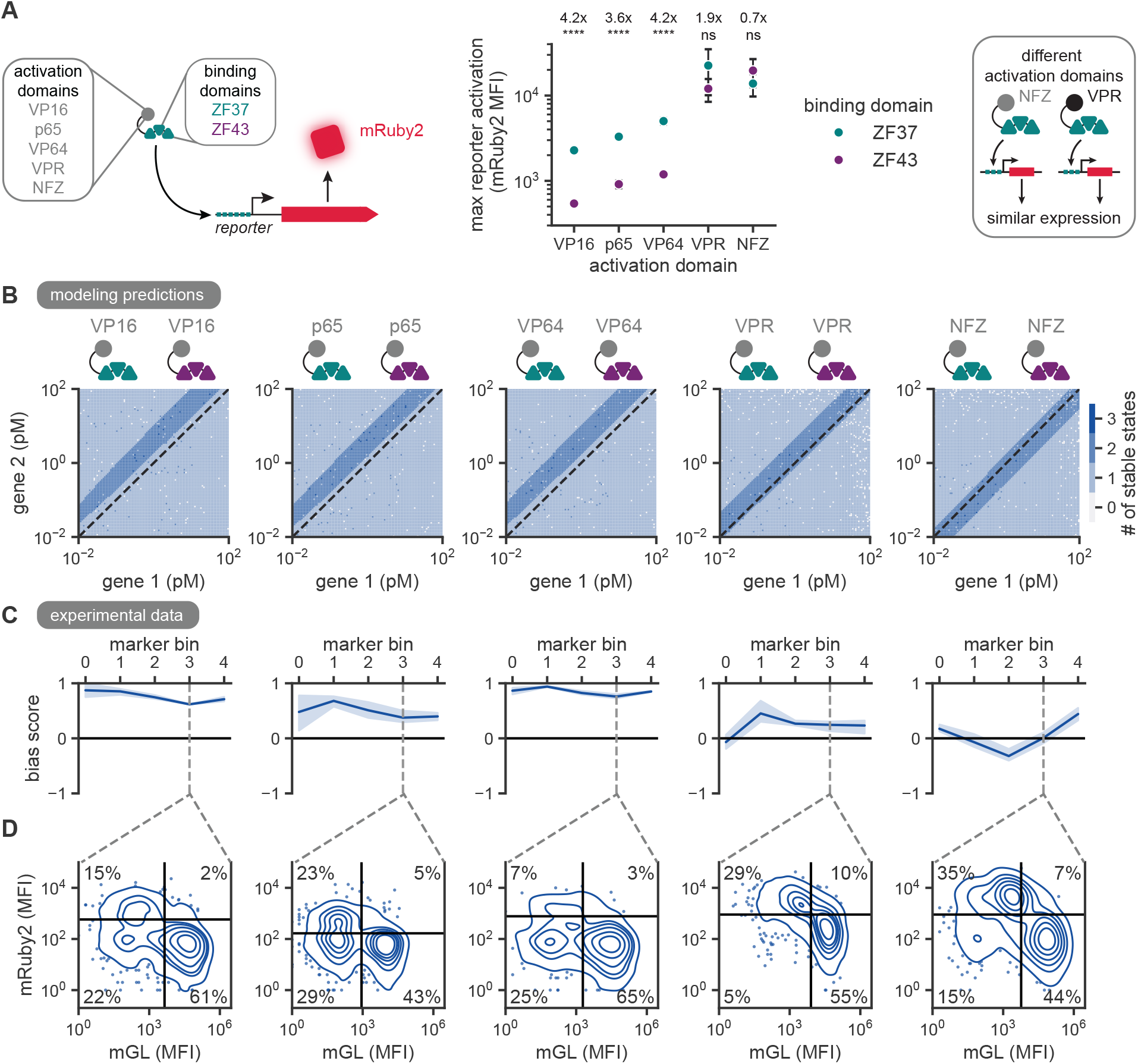
Circuit bias can be reduced by balancing transcriptional activation strength. (**A**) Maximum reporter activation with varying activation domain and binding domain. The activation domains tested here include VP16, p65, VP64, VPR, and NFZ. Each was paired with the ZF37 and ZF43 binding domains and characterized in HEK293T cells via poly-transfection. Binned data from Figure S6 was fit to a Hill function with a Hill coefficient of one. Fits are displayed in Figure S7. Points represent the maximum activation parameter obtained by fitting data from at least three biological replicates, and error bars represent the standard deviation of the fitted parameter. Annotations represent the ratio between parameter values for the ZF37 and ZF43 domains. These values are reported in Table S2. ****: *p <* 0.0001, n.s.: *p >* 0.05, two-sided Z-test [26]. (B) Deterministic modeling of the circuit with varying activation domains over different concentrations of gene 1 and gene 2. Parameters used for activation strength of each domain are reported in Table S2. Unless otherwise stated, parameters used in modeling are reported in Table S1. The deterministic model is defined by Equations S1 to S10 as derived in the supplementary text. Dashed line indicates 1:1 copy number ratio of the two genes. (**C**) Plot of gene circuit bias for poly-transfection bins with proportional levels of both genes. Data points represent the mean of three biological replicates, and shaded regions represent the 95% confidence interval. Grey dashed line indicates the selected bin shown in **D**. VP16 circuit data displayed in the furthest left plot was previously reported in Figure 3B. (**D**) Density plots depicting one selected poly-transfection bin indicated in the above bias plot. Data is shown for one representative biological replicate. Points represent the 5% of probability mass lying outside the lowest iso-proportion contour. Data is in arbitrary units from a flow cytometer. Density plots for all bins are displayed in Figures S4, S8, S9, S10 and S11. VP16 circuit data displayed in the furthest left plot was previously reported in Figure 3C. Black lines indicate the quadrants used in calculating the bias score. Quadrant gates are set based on expression distributions in tagBFP bin 0 and iRFP670 bin 0, which are reported in Figures S4, S8, S9, S10 and S11.

Using parameters scaled based on experimentally determined reporter activation levels (Table S2), we modeled the circuit stability with varying activation domains. Again, these mathematically stable attractor states correspond to two distinct cellular populations; however, this modeling does not incorporate biological noise or cell cycle, which would be necessary to predict long-term stability of the states. With the imbalanced VP16, p65, or VP64 activators, modeling predictions indicate that two distinct attractor states are only achieved with higher copy numbers of gene 2 compared to gene 1 (Figure 4B). The predicted bistable parameter regime does not include any conditions where gene 1 and gene 2 are present at equal copy numbers. In comparison, modeling the circuit with parameters scaled to the NFZ or VPR experimental characterization data indicates that the system has two states when the genes are at equal copy numbers (Figure 4B).

Performing poly-transfection characterization for each of these activation domain pairings (Figures S8, S9, S10 and S11), we considered the circuit bias across the diagonal bins, as previously done for VP16 (Figure 3B). The NFZ and VPR domains each yield similar activation when paired with either the ZF37 or ZF43 domains (Figure 4A), and the circuits using these domains yield more balanced induction of both state 1 and state 2 at equal copy numbers of each gene (Figure 4C). In comparison, the p65 and VP64 domains induce higher activation when paired with ZF37 (Figure 4A), causing the DACs using these domains to exhibit a bias towards state 1, similar to the VP16-based design (Figure 4C). Looking again at the distributions of mGL and mRuby2 expression within one of these bins, we find that there is robust induction of state 2 only for the VPR and NFZ circuits (Figure 4D). Despite the relatively unbiased distributions of the VPR-based circuit, the two states are close in expression level, with above-background expression of the opposite gene in both states (Figure S10). This is potentially due to the poor repression of VPR-based transcriptional activation by the ZF repressors used here (Figure S6B). Comparing across all activation domains, the NFZ domain enables the most unbiased state specification at one-to-one copy numbers (Figure 4D). Additionally, the NFZ domain (∼ 0.5 kb) is significantly smaller in cargo size than the VPR domain (∼ 1.5 kb) and is derived from human sequences, potentially increasing delivery efficiency and reducing immunogenicity relative to VPR. In alignment with mathematical modeling of the circuit at equal copy numbers, substitution of the VP16 activation domains for NFZ domains promotes more robust induction of both DAC states by balancing transcriptional activation.

### Competition for transcriptional co-activators reinforces the two attractor states

To test whether activation strength alone predicts circuit performance, we quantified circuit performance when pairing different activation domains that have similar strengths. If transcriptional activation strength alone determines the bias of the circuit, we expect that mismatched activation domains would perform well as long as the activators generate comparable reporter expression. To probe this effect, we specifically considered circuits with varying pairings of NFZ and VPR-based activators. These domains yield similar levels of reporter activation but rely on distinct sets of transcriptional co-activator species (Figure 4A) [27, 28]. We performed poly-transfection of each pairing (Figures S10, S11, S12 and S13) and observed the resulting expression distributions in a bin with high copy number of both genes (Figure 5A). For conditions with matched activation domains, the circuit maintains a two-state expression distribution even at high copy number (Figure 5A). In contrast, the conditions with mismatched activation domains display more bias or a single state (Figure 5A). In both cases, the circuit is biased towards the state specified by the VPR-based activator (Figure 5A) despite the NFZ domain inducing similar reporter activation in previous characterization experiments (Figures 4A and S6). Therefore, transcriptional activation strength alone is not a good predictor of circuit performance. This indicates that there are other phenomena that are currently neglected by our DAC model that limit its predictive capacity.

**Figure 5.**
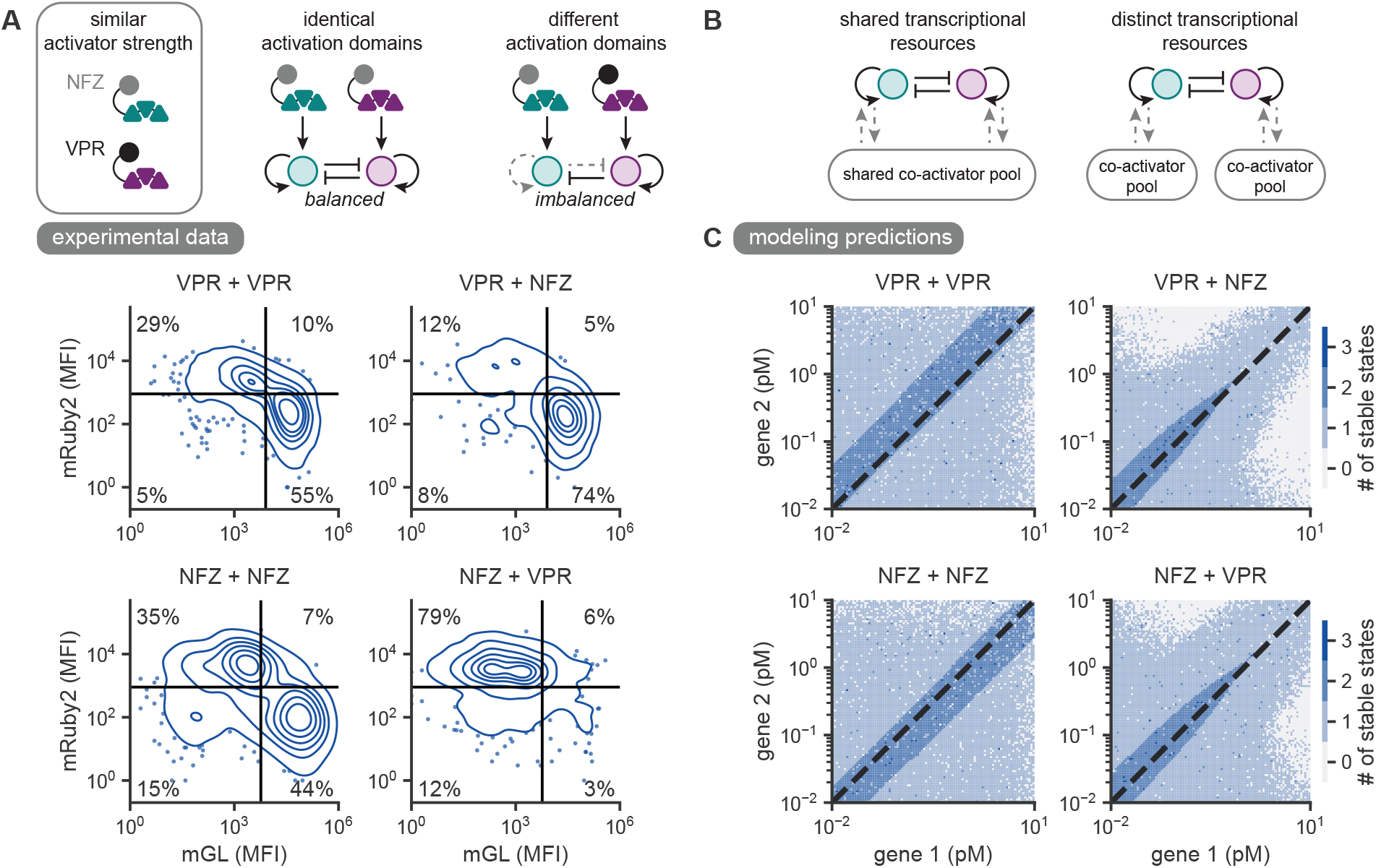
Competition for transcriptional co-activators reinforces the two attractor states. (**A**) Matched activation domains maintain two attractor states more robustly than mismatched activation domains at high copy numbers. Poly-transfection of the circuit with varying activation domains in HEK293T cells. Data is shown for bin 3 of each transfection marker, and data for all bins is reported in Figures S10, S11, S12 and S13. Density plots depict one representative biological replicate. Points represent the 5% of probability mass lying outside the lowest iso-proportion contour. Data is in arbitrary units from a flow cytometer. Quadrant gates are set based on expression distributions in tagBFP bin 0 and iRFP670 bin 0, which are reported in Figures S10, S11, S12 and S13. (**B**) At high copy numbers, identical activation domains share a pool of transcriptional co-activator resources. Different activation domains may utilize distinct pools of transcriptional resources. Choice of activation domain pairings impacts coupling between the two genes due to resource competition. (**C**) Competition for transcriptional co-activators is predicted to reinforce the two attractor states at high copy numbers. Deterministic modeling of the circuit with varying activation domains and shared transcriptional resources over varying concentrations of gene 1 and gene 2. Parameters used in modeling are reported in Tables S1 and S2. The deterministic model is defined by Equations S3 to S12 as derived in the supplementary text. Dashed line indicates 1:1 copy number ratio of the two genes.

Resource competition can significantly impact the performance of gene circuits at high expression levels [29–31]. Dual-attractor gene networks, in particular, can be impacted significantly by the sharing of transcriptional or translational resources [32]. This is relevant for the selection of activation domains that enable robust circuit performance across copy number regimes. Depending on the identity of the activation domain, different co-activator species are recruited to initiate transcription. For a system with two matching activation domains, both genes rely on the same pool of transcriptional resources, further coupling the two genes (Figure 5B). Therefore, we sought to understand how sharing of transcriptional resources can impact the distributions of the two attractor states.

To probe the effects of transcriptional resource limitations, we modified the ODE model of the circuit to account for limited transcriptional resources (Equations S11 and S12). In the case of circuits containing both NFZ and VPR domains, the two genes utilize separate transcriptional resource pools (Figure 5B). As copy number increases, the available co-activator pools become more depleted, resulting in a reduction in transcriptional activation. However, since the resource pools for different activation domains are not shared, this resource competition can have unbalanced effects on the two genes. In the modeling predictions, this causes the system to collapse to a single state at high copy numbers (Figure 5C). In the case of circuits containing either matched NFZ or matched VPR activation domains, the depletion of the available co-activator pool equally impacts both of the genes. This is predicted to act as an additional form of mutual inhibition between the two genes that maintains two attractor states even in a high-copy, resource-limited regime (Figure 5C). Overall, the results of this deterministic modeling and poly-transfection characterization suggest that activation domain strength alone is not sufficient to predict DAC performance. Instead, selection of matched activation domains may contribute to more robust maintenance of two states at high copy numbers through transcriptional resource competition. The depletion of shared resources can act as an additional form of mutual inhibition, allowing the circuit to maintain two states in a regime where it would otherwise collapse to a single state.

### Layering microRNA-based mutual repression can reduce circuit bias

Native circuits use multiple modes of regulation to reinforce cell state. A central challenge in engineering single-cargo DACs is introducing sufficient regulatory control without substantially increasing genetic cargo size. We wondered if layering microRNA-based repression on top of the existing biochemical regulation could improve DAC performance [33]. MicroRNA-based repression can be implemented independently from transcriptional repression and is compact, adding approximately 400 base pairs to each gene. This approach was originally applied to a circuit developed by Kramer *et al* by encoding synthetic introns containing microRNA sequences complementary to the opposite gene [11, 33]. Here, we investigated microRNA-based repression as a tool to reduce circuit bias when using imbalanced transcriptional activators. Upon expression of one of the genes, the synthetic microRNA will be spliced and processed, allowing it to hybridize with a target site encoded in the 3’ UTR of the opposite gene’s transcript (Figure 6A). The microRNA binding can inhibit translation and induce transcript degradation. When modeled with imbalanced VP16 activation, the additional microRNA-based repression expands the bistable parameter regime compared to the original circuit (Figures 4B and 6B). Under these conditions, the circuit is bistable at some 1:1 copy numbers, although the model predicts monostability at low copy numbers (Figure 6B). Therefore, additional mutual inhibition encoded by microRNAs may promote less biased adoption of both states when activation strength cannot be effectively balanced.

**Figure 6.**
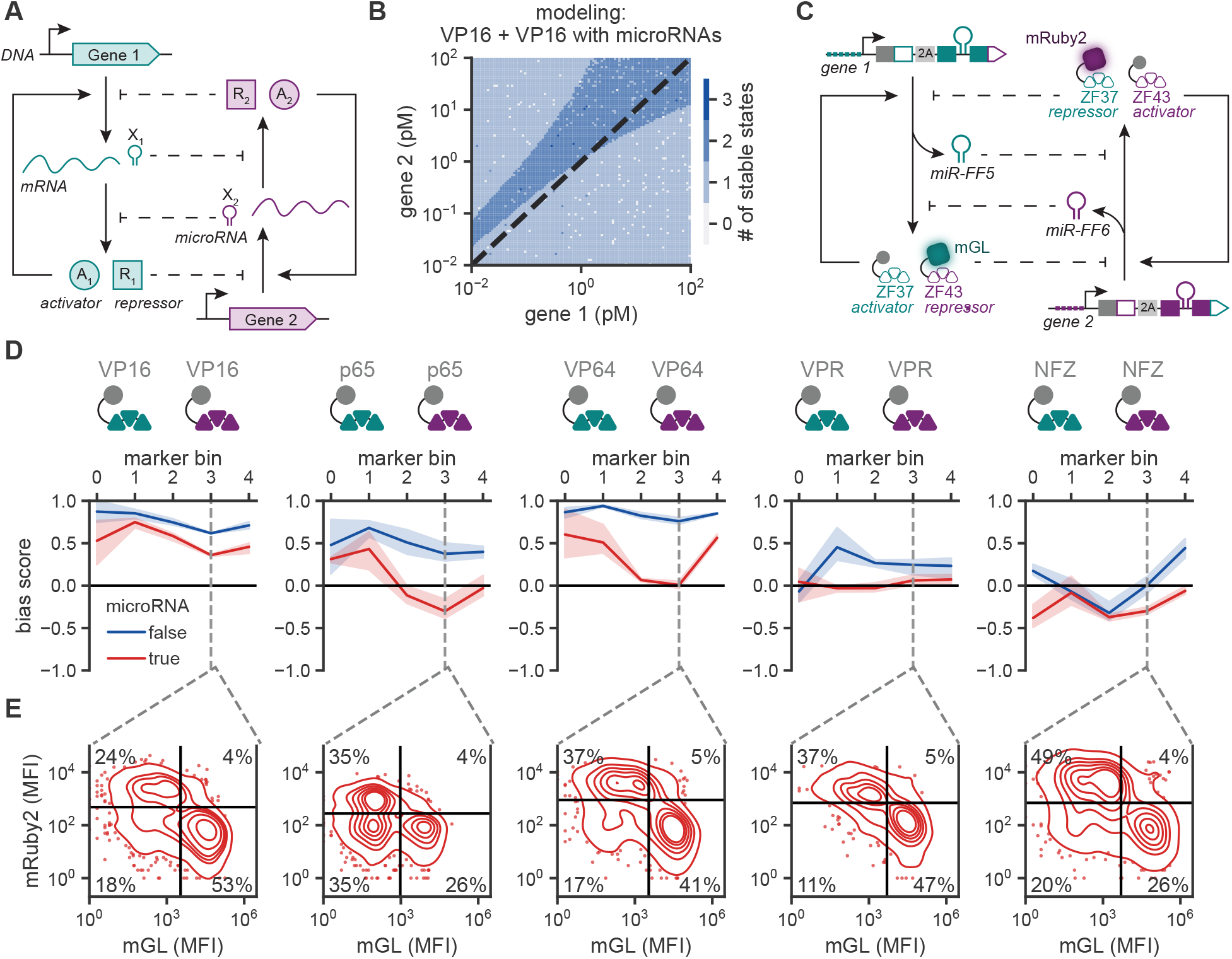
Layering microRNA-based mutual repression can reduce circuit bias. (**A**) Transcriptional repression and microRNA-based repression can be combined for additional control. Here, each gene encodes for a transcription activator, a transcriptional repressor, and a microRNA. Each activator can bind to its own promoter, implementing positive feedback. Each repressor can bind to the promoter of the opposite gene, inhibiting transcriptional activation. Each microRNA can bind to the mRNA transcript of the opposite gene, inhibiting translation. Both methods implement mutual repression (dashed lines). (**B**) Deterministic modeling of the circuit with two VP16 activation domains (*β*_1_ = 6 and *β*_2_ = 1.4) and microRNA-mediated repression. Unless otherwise stated, parameters used in modeling are reported in Table S1. The deterministic model is defined by Equations S1 to S10 as derived in the supplementary text. Dashed line indicates 1:1 copy number ratio of the two genes. The bistable parameter regime overlaps the 1:1 diagonal at medium-to-high copy numbers. (**C**) microRNA-based repression is added to the previous DAC design by embedding microRNA-containing synthetic introns within the coding sequences of the fluorescent proteins. The synthetic intron for gene 1 is embedded within the mGL coding sequence and encodes for the miR-30 backbone and a miR-FF5 hairpin. The synthetic intron for gene 2 is embedded within the mRuby2 coding sequence and encodes for the miR-30 backbone and a miR-FF6 hairpin. (**D**) Plot of gene circuit bias for poly-transfection bins with proportional levels of both genes. Data points represent the mean of three biological replicates, and shaded regions represent the 95% confidence interval. Grey dashed line indicates the selected bin shown in **E**. Circuit data without additional microRNA repression was previously reported in Figure 4C. (**E**) Density plots depicting one selected poly-transfection bin indicated in the above bias plot. Data is shown for one representative biological replicate. Points represent the 5% of probability mass lying outside the lowest iso-proportion contour. Data is in arbitrary units from a flow cytometer. Density plots for all bins are displayed in Figures S15, S16, S17, S18 and S19. Black lines indicate the quadrants used in calculating the bias score. Quadrant gates are set based on expression distributions in tagBFP bin 0 and iRFP670 bin 0, which are reported in Figures S15, S16, S17, S18 and S19.

To apply this additional microRNA-based control within our circuit, miR-FF5 and miR-FF6 can be encoded within the fluorescent protein domains of the transcriptional repressors (Figure 6C). We selected the miR-FF5 and miR-FF6 sequences since they are orthogonal to native human transcript sequences and are similar in knockdown strength [34–36]. When encoded as a synthetic intron within a fluorescent protein, these microRNAs enact a dose-dependent reduction in reporter expression for transcripts containing their cognate target site (Figure S14). Characterizing this circuit architecture in poly-transfection, we find that it reduces the bias of the circuit towards state 1 compared to the circuit without additional microRNA-based mutual inhibition (Figure 6D). However, the microRNA-based repression has the largest effect at intermediate copy numbers, potentially indicating that the RNA-induced silencing complex (RISC) is being saturated at high copy numbers [36]. This effect generalizes across all activation domains except NFZ (Figures S16, S17, S18 and S19), for which the circuit is already well-balanced. Looking at the distribution of mGL and mRuby2 expression within one bin with high expression of both marker plasmids, we observe more robust induction of state 2 even in conditions with significantly different activation strengths (Figure 6E). Therefore, additional microRNA-mediated repression can reduce bias due to an imbalance in transcriptional activation strength by encoding additional mutual repression between the genes.

### Optimized single-cargo, dual-attractor circuits enable robust induction of two cell states

Selection of transcriptional activation domains and addition of microRNA-mediated repression allowed us to achieve virtually unbiased cell-state commitment with co-transfection of two plasmids (Figure 6). However, delivery of two separate cargoes is still a barrier to translation of this DAC circuit beyond transient transfection [17]. With the reduced cargo size of the ZF and NFZ domains, the DAC is compact enough to be encoded on a single plasmid. By ensuring that each gene is present at a one-to-one copy number, a single-cargo design should enable more homogeneous performance of the DAC across polyclonal populations [17]. However, co-localization of both DAC genes on a single plasmid also introduces biophysical coupling that can impact the balance of the DAC states [19, 21]. Gene syntax dictates the sign and degree of transcriptional coupling between adjacent genes [20]. Therefore, we tested the best-performing DAC circuit in each of the possible two-gene syntaxes (i.e. divergent, convergent, tandem 1, tandem 2) (Figure 7A). We found that the tandem syntax with gene 2 in the upstream position achieved significantly better balance than all other syntaxes, with the lowest bias score (Figure S20) and the highest ratio of state 2 cells relative to state 1 cells (Figure 7B). Optimal balance from the tandem syntax for this mammalian circuit contrasts with previous experimental work in bacteria [21] and biophysical modeling predictions [19], which found that the convergent syntax is the best performing. Differences in the predictions and experimental results may reflect a remaining imbalance in the circuit properties such as differences the kinetics of transcription, which the tandem syntax may mitigate.

**Figure 7.**
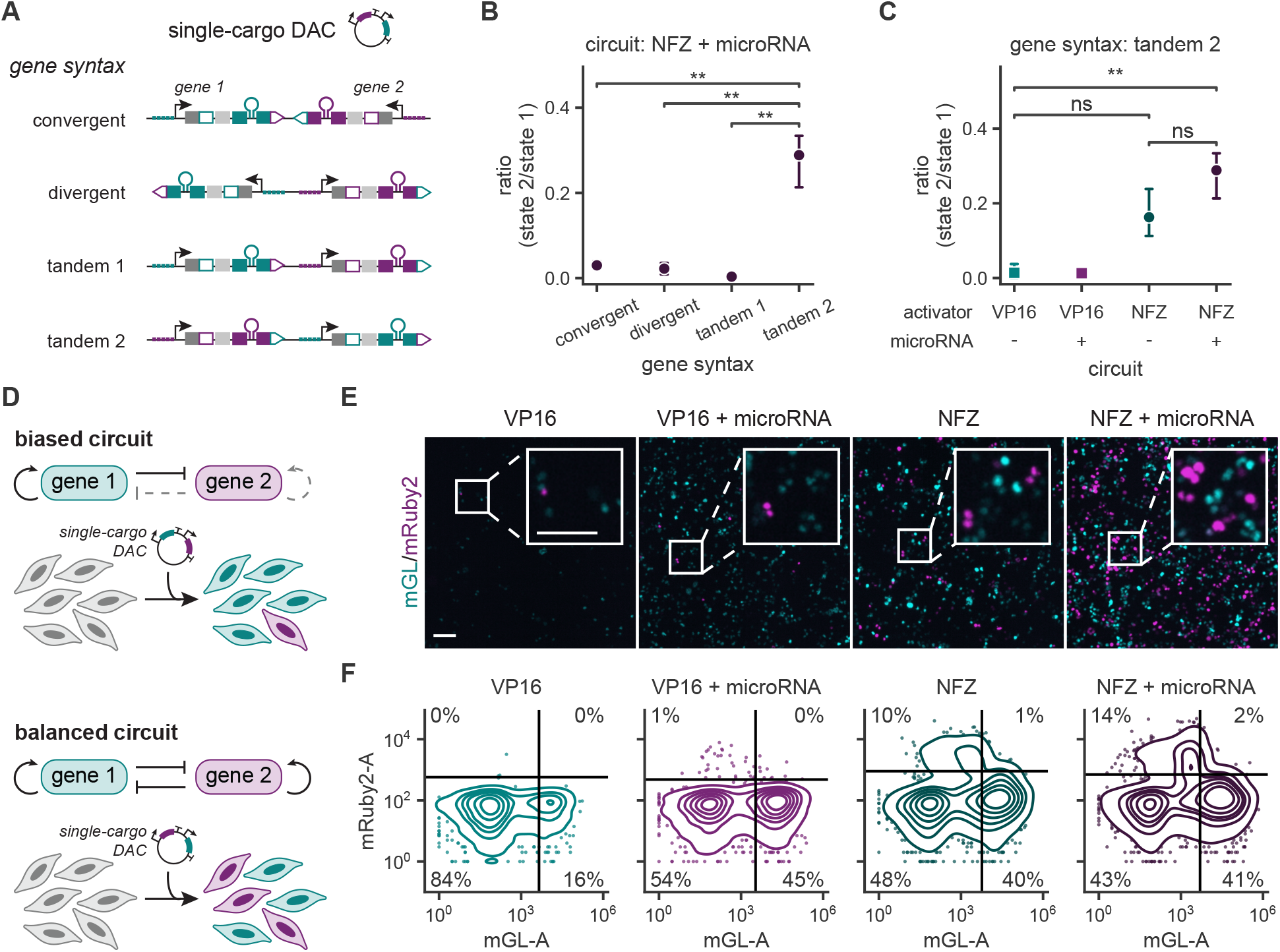
Optimized single-cargo, dual-attractor circuits enable robust induction of two cell states. (**A**) The compact nature of this DAC design allows both circuit genes to be encoded on a single plasmid, ensuring that both genes are delivered at the same copy number. However, co-localization of genes introduces biophysical coupling, which is modulated by gene syntax [19, 21]. HEK293T cells were transfected with varying single-cargo DACs, and expression of circuit genes was quantified by flow cytometry at 2 dpt. (**B**) Ratio of cells in state 2 (mRuby2+) to cells in state 1 (mGL+) for varying gene syntax with the NFZ + microRNA circuit. Data points represent the mean of three biological replicates, and error bars represent the 95% confidence interval. **: 0.01 ≥ *p >* 0.001, two-sided t-test with Bonferroni correction. (**C**) Ratio of cells in state 2 (mRuby2+) to cells in state 1 (mGL+) for varying DAC design. All circuits were encoded in the tandem orientation with gene 2 upstream. Data points represent the mean of three biological replicates, and error bars represent the 95% confidence interval. ns: *p >* 0.05, **: 0.01 ≥ *p >* 0.001, two-sided t-test with Bonferroni correction. (**D**) Through a combination of biochemical and biophysical tuning, the bias of single-cargo DACs can be reduced to achieve a more balanced ratio of the two attractor states. (**E**) Images of HEK293T cells transfected with varying single-cargo DAC designs in the tandem syntax with gene 2 upstream. Scale bars represent 75 µm. (**F**) Density plots depicting expression for DAC designs in the tandem syntax with gene 2 upstream. Data is shown for one representative biological replicate. Points represent the 5% of probability mass lying outside the lowest iso-proportion contour. Data is in arbitrary units from a flow cytometer. Black lines indicate the quadrants used in calculating the state ratios. Quadrant gates are set based on expression distributions in tagBFP bin 0 and iRFP670 bin 0 from poly-transfection experiments of the corresponding two-cargo circuits, which are reported in Figures S4, S11, S15 and S19.

We also tested previous DAC designs in the tandem 2 gene syntax to investigate whether the biochemical optimization in a co-transfection context generalized to single-cargo designs. In agreement with prior results, the combination of DAC activation domain selection and addition of supplemental microRNA repression acts synergistically to reduce circuit bias (Figure S20) and improve the balance of state 2 cells relative to state 1 cells (Figure 7C). In the single-cargo context, both biochemical and biophysical optimization are required to achieve robust circuit performance (Figure 7D). The single-cargo design (Figure 7) shows greater bias compared to the two-cargo design (Figure 6) but eliminates copy number as a source of variance. Together, we show that combined biophysical and biochemical tuning methods can enable robust induction of both DAC states from a single cargo (Figures 7E and 7F), laying the foundation for translation of DACs to application-relevant cell types.

## Discussion

Engineering single-cargo DACs requires strategies that improve circuit performance without substantially increasing the genetic footprint of the circuit. Here, we establish model-guided design principles for engineering compact, single-cargo DACs that generate two states. Starting from a previously developed TALE-based circuit that required four cargoes and more than 15 kilobases of circuit sequence, we replaced the large DNA-binding domains with compact zinc-fingers, reducing the circuit footprint to under 5 kilobases. Through combined biochemical and biophysical tuning, we consolidated the circuit into a single genetic cargo of less than 11 kilobases while reducing the initial state bias from 5:1 to approximately 3:1. Importantly, this circuit provides the controllability and deliverability of a single cargo (Figure 1E).

In developing the single-cargo DAC, we find that balance in transcriptional activation strength is key to achieving unbiased state induction. The same transcriptional activation domains can have significantly different reporter induction depending on the paired binding domain, suggesting multiple modes of tuning circuit activity (Figure 4). While we identify a pairing of transcriptional activators that have similar strength to reduce circuit bias (Figure 4), this design space is limited and discrete in nature. Optimization through varying activation or binding domains requires screening large numbers of circuit designs. Improved programmability requires methods of tuning transcriptional activation in a more continuous manner. Potentially, transcriptional activation can be tuned by modifying the affinity of DNA binding domains [13, 25] or the distance between TF binding sites and the minimal promoter in the promoter design [25, 37]. Multiple studies have demonstrated that arginine-to-alanine mutations in the binding domain of a ZF can reduce binding affinity, with more mutations corresponding to lower reporter activation levels [13, 25, 38]. Additionally, increasing the distance between TF binding sites and a minimal promoter decreases reporter activation [25, 37]. Both of these approaches offer more predictable and continuous tuning of transcriptional activation. However, these techniques primarily reduce expression, so systems with high activity are required as a starting point.

In testing pairings of activation domains that have similar strengths but different transcriptional co-activators, we found activation strength alone is not sufficient to predict DAC performance (Figure 5). The identity of the activation domain itself plays a significant role in circuit performance, and resource competition potentially shapes stability at high copy numbers (Figure 5). Despite having similar strengths, circuits using the NFZ and VPR activation domains can only robustly induce two states when the activation domains are matched. In the high copy number context of transfection, competition for shared transcriptional co-activators can act as a form of mutual inhibition, layering on top of the programmed biochemical regulation. This effect may be generalizable to other two-state attractor circuits using transcriptional activation [12, 13]. However, for two-state systems relying only on cooperative transcriptional repression [11], resource competition is predicted to reduce performance, potentially causing the circuit to be monostable [39, 40]. More work is needed to understand how these resource competition effects impact circuits with different layers and signs of biochemical regulation.

As single-cargo DACs require strategies for layering regulatory control without substantially expanding the genetic cargo, we applied a layer of microRNA-mediated mutual repression to reduce circuit bias (Figure 6). Synthetic microRNA-based regulation functions reliably in low-copy contexts and in primary cells [36, 41], making it a useful tool as bipotent circuit development moves closer to translation. By adding an orthogonal biochemical repression method, we reduced the circuit bias by reinforcing the mutual inhibition central to cell-state commitment. The degree of bias mitigation from microRNA-mediated mutual repression is smaller in magnitude than the impact of activator substitution, however, the range of stability is wider using the microRNA. To achieve a synergistic reduction in bias, activator optimization and microRNAs can be combined as complementary fine-tuning approaches.

Finally, testing these DAC designs in a single-cargo context, we find that gene syntax is a key determinant of performance (Figure 7). Gene syntax modulates the biophysical coupling between DAC genes, adding another layer of regulation for tuning circuit activity [19, 21]. We find that placing the weaker gene in the upstream position in tandem syntax yields the most balanced cell-state commitment. These findings establish gene syntax as a complementary design variable that can be used alongside biochemical regulation to optimize circuit behavior under strict cargo-size constraints. As the impact of biophysical coupling varies between plasmid-based expression and genomic integration, the optimal arrangement will need to be reevaluated in integrated systems [19, 20].

Overall, this work establishes multiple biochemical regulation strategies for reducing bias of a dual-attractor circuit. Importantly, these approaches do not rely on copy number variation and significantly minimize the size and number of the genetic cargoes require to generate dual states. As transient delivery limits our ability to assess long term stability of the DAC states, future work is needed to explore the stability of DAC-generated states. This work provides a critical step towards development for stable circuit performance across a polyclonal population. Together, these advances establish a compact, single-cargo DAC design that lays the foundation for stable circuit performance across polyclonal cell populations, enabling future applications of circuit-regulated cell and gene therapies.

## Data and Materials Availability

Raw flow cytometry data and fluorescence images will be deposited at Zenodo. Plasmids listed in Table S3 will be deposited at Addgene. Code for data analysis and mathematical modeling will be made available at https://github.com/GallowayLabMIT.

## Acknowledgments

We thank Maëlle Jacquemettaz, Albert Blanch-Asensio, Kasey S. Love, Amy Oh, and Maria F. Castellanos for their generous experimental support. Additionally, we thank Brittany A. Lende-Dorn, Kasey S. Love, and Mary E. Ehmann for their valuable feedback on this manuscript.

## Author Contributions

E.L.P: Conceptualization; Data curation; Formal analysis; Investigation; Methodology; Project Administration; Visualization; Writing - original draft; Writing - review & editing. N.N: Conceptualization; Formal analysis; Investigation; Methodology; Writing - original draft; Writing - review & editing. P.M.N: Formal analysis; Investigation; Writing - review & editing. D.R.G: Formal analysis; Investigation; Writing - review & editing. D.D.V: Conceptualization; Funding acquisition; Project administration, Supervision; Writing - review & editing. K.E.G: Conceptualization; Funding acquisition; Project administration, Supervision; Writing - original draft; Writing - review & editing.

