## Supplement for "Model-guided design of single-cargo, dual-attractor gene circuits"

### Supplementary Information

#### Materials and Methods

##### ODE simulations

The reaction networks derived in the supplementary text were simulated in Julia using the `Catalyst.jl` package [42]. Parameters used in the model are reported in Table S1. Steady states for each set of parameters were computed using a homotopy continuation algorithm implemented through the `HomotopyContinuation.jl` package [43]. The stability of each steady state was computed using a built-in function that considers the Jacobian of the ODE system.

##### Cloning

Single-gene plasmids were assembled using a multi-level Golden Gate cloning scheme. Individual genetic parts were amplified from commercial DNA sources and inserted into a corresponding part positioning vector (pPV) backbone via Gibson Assembly using Hifi DNA Assembly Master Mix (NEB, M5520). ZF-responsive promoters, ZF binding domains, and the VP16, VP64, and VPR activation domains were amplified from plasmids deposited by Donahue *et al* [25]. The p65 activation domain was amplified from a plasmid deposited by Li *et al* [38]. The NFZ activation domain was amplified from a plasmid deposited by Tycko *et al* [28]. MicroRNA sequences were assembled based on sequences reported by Leisner *et al* [35]. Full transcriptional units were assembled in a BsaI (NEB, R3733L) Golden Gate reaction to yield the kanamycin-resistant plasmids (pShips) used in transfection experiments. Two-gene plasmids for delivery of DAC circuits as a single cargo were assembled in a PaqCI (NEB, R0745L) Golden Gate reaction. Plasmids used in transfection experiments are reported in Table S3.

##### Cell culture

HEK293T cells (ATCC, CRL-3216) were cultured using DMEM (Genesee Scientific, 25-501) plus 10% FBS (Genesee Scientific, 25-514H) and incubated at 37°C with 5% CO2. Cells were passaged at 80-90% confluence, in which spent media was aspirated, and cells were washed with PBS (Sigma-Aldrich, P4417-100TAB) then subsequently dissociated with 0.25% Trypsin-EDTA (Genesee Scientific, 25-510) diluted in PBS. After four minutes, cells were spun down at 400 rcf for 5 minutes, resuspended in media, then transferred to a new flask. Media was replaced with fresh DMEM + 10% FBS every 2-3 days.

##### Plasmid transfection

In preparation for experiments, HEK293T cells were counted using a hemocytometer, seeded with 0.1% gelatin coating (Sigma-Aldrich, G1890-100G) at a density of 30,000 cells per 96-well or 150,000 cells per 24-well, and transfected 24 hours later. Transfection mixes were prepared using a ratio of 4 µg PEI to 1 µg DNA. First, a master mix of PEI and KnockOut™ DMEM (ThermoFisher Scientific, 10-829-018) was prepared and incubated for a minimum of 10 minutes. This master mix was then added to DNA mixes. These condition mixes were further incubated for 10 to 15 minutes and then added on top of the growth media in the plate. Media was replaced with fresh DMEM + 10% FBS after 24 hours. At 2 days post transfection, HEK293T cells were dissociated by adding 0.25% Trypsin-EDTA (diluted in PBS) for four minutes, followed by quenching with an equal volume of DMEM + 10% FBS. After centrifuging at 500 rcf for 5 minutes, cells were resuspended in PBS and transferred to a v-bottom plate for flow cytometry.

*Co-transfection of initial circuit design (Figure 2)*

Experiments were performed at 96-well scale, and one co-transfection mix was prepared per condition. Co-transfection mixes were prepared containing 38 ng of gene 1 plasmid, 38 ng of gene 2 plasmid, and 38 ng of transfection marker plasmid per 96-well.

*Poly-transfection for repressor characterization (Figures S2 and S6B)*

Poly-transfection was performed by separately preparing co-transfection mixes and adding each co-transfection mix on top of the growth media in the plate. Experiments were performed at 24-well scale, and three separate co-transfection mixes were prepared per condition: (1) 225 ng of reporter plasmid and 225 ng of transfection marker plasmid per 24-well, (2) 450 ng of repressor plasmid per 24-well, and (3) 113 ng of activator plasmid and 113 ng of filler plasmid per 24-well.

*Poly-transfection for activator characterization (Figures 4A and S6A)*

Poly-transfection was performed by separately preparing co-transfection mixes and adding each co-transfection mix on top of the growth media in the plate. Experiments were performed at 96-well scale, and two separate co-transfection mixes were prepared per condition: (1) 56 ng of reporter plasmid and 56 ng of transfection marker plasmid per 96-well and (2) 28 ng of activator plasmid and 28 ng of filler plasmid per 96-well.

*Poly-transfection for microRNA characterization (Figure S14)*

Poly-transfection was performed by separately preparing co-transfection mixes and adding each co-transfection mix on top of the growth media in the plate. Experiments were performed at 96-well scale, and two separate co-transfection mixes were prepared per condition: (1) 56 ng of reporter plasmid and 56 ng of transfection marker plasmid per 96-well and (2) 56 ng of microRNA plasmid and 56 ng of filler plasmid per 96-well.

*Poly-transfection for DAC bias characterization (Figures 3, 4, 5 and 6)*

Poly-transfection was performed by separately preparing co-transfection mixes and adding each co-transfection mix on top of the growth media in the plate. Experiments were performed at 96-well scale, and two separate co-transfection mixes were prepared per condition: (1) 56 ng of gene 1 plasmid and 56 ng of marker 1 plasmid per 96-well and (2) 56 ng of gene 2 plasmid and 56 ng of marker 2 plasmid per 96-well.

*Co-transfection of single-cargo DAC designs (Figure 7)*

Experiments were performed at 96-well scale, and one co-transfection mix was prepared per condition. Co-transfection mixes were prepared containing 76 ng of DAC plasmid and 38 ng of transfection marker plasmid per 96-well.

#### Flow cytometry

All flow cytometry data was collected using an Attune NxT flow cytometer with channel mappings and voltages reported in Table S4. Single cells were selected using forward scatter and side scatter gates. Transfected cells were gated based on expression of a co-transfected marker. For poly-transfection analysis, cells were separated into equal-width or equal-quantile bins where noted in figure legends. For each circuit design, the mGL gate was set based on mGL expression within the lowest tagBFP bin, and the mRuby2 gate was set gased on mRuby2 expression within the lowest iRFP670 bin.

#### Imaging

Cells were imaged in PBS or FluoroBrite™ DMEM (ThermoFisher Scientific, A1896701) to minimize background fluorescence. Images were captured using a Nikon Ti-2E epifluorescence microscope at 4x magnification.

### Supplementary Text

#### Model derivation

Here, we derive a deterministic model for dual-attractor gene circuits utilizing positive transcriptional feedback and mutual repression via transcriptional repressors and microRNAs. We start by considering the equations related to expression of Gene 1.

*Transcription*

Transcription of Gene 1 is governed by the following reactions:

$$G_1 \xrightarrow{\alpha_1} G_1 + T_1$$

$$A_1 + G_1 \underset{k_{A1,unbind}}{\overset{k_{A1,bind}}{\longleftrightarrow}} C_{A1,G1} \xrightarrow{\beta_1} C_{A1,G1} + T_1$$

$$R_2 + G_1 \underset{k_{R2,unbind}}{\overset{k_{R2,bind}}{\longleftrightarrow}} C_{R2,G1}$$

where the unbound DNA encoding Gene 1 is $G_1$, the primary RNA transcript is $T_1$, the transcriptional activator encoded by Gene 1 is $A_1$, the transcriptional repressor encoded by Gene 2 is $R_2$, and the complexes of each TF with the DNA are $C_{A1,G1}$ and $C_{R2,G1}$, respectively. Similarly, $\alpha_1$ is the rate constant for basal transcription, $\beta_1$ is the rate constant for activated transcription, $k_{A1,bind}$ and $k_{A1,unbind}$ are the rate constants for activator association and dissociation, and $k_{R2,bind}$ and $k_{R2,unbind}$ are the rate constants for repressor association and dissociation. The rate of transcription of Gene 1 is:

$$f_{1,txn} = \alpha_1 G_1 + \beta_1 C_{A1,G1}$$

Assuming that the dynamics of TF binding are fast compared to transcription, we can solve for the concentrations of the complexes and write a conservation equation for $G_1$:

$$C_{A1,G1} = \frac{A_1 G_1}{K_{D,A1}} \qquad C_{R2,G1} = \frac{R_2 G_1}{K_{D,R2}}$$

$$G_{1,tot} = G_1 + C_{A1,G1} + C_{R2,G1}$$

$$G_1 = \frac{G_{1,tot}}{1 + \frac{A_1}{K_{D,A1}} + \frac{R_2}{k_{D,R1}}}$$

where $K_{D,A1}$ is the dissociation constant of the activator, $K_{D,R2}$ is the dissociation constant of the repressor, and $G_{1,tot}$ is the total copy number of Gene 1. With these simplifications, the transcription rate of Gene 1 is written as a function of $A_1$ and $R_2$:

$$f_{1,txn}(A_1, R_2) = G_{1,tot} \frac{\alpha_1 + \frac{\beta_1 A_1}{K_{D,A1}}}{1 + \frac{A_1}{K_{D,A1}} + \frac{R_2}{K_{D,R2}}}$$

Similarly, the transcription rate of Gene 2 is expressed as a function of $A_2$ and $R_1$:

$$f_{2,txn}(A_2, R_1) = G_{2,tot} \frac{\alpha_2 + \frac{\beta_2 A_2}{K_{D,A2}}}{1 + \frac{A_2}{K_{D,A2}} + \frac{R_1}{K_{D,R1}}}$$

*Splicing*

Splicing of the primary RNA transcript for Gene 1 is represented by the following reaction:

$$T_1 \xrightarrow{\sigma} M_1 + x\ X_1$$

$$x = \begin{cases} 0, & \text{microRNA-containing intron not present} \\ 1, & \text{microRNA-containing intron present} \end{cases}$$

where $M_1$ is the mature mRNA transcript and $X_1$ is the spliced microRNA. The parameter $\sigma$ represents the splicing rate constant, and $x$ is the stoichiometric coefficient of the microRNA determined by the presence or absence of a microRNA-containing intron in the gene [44]. We assume the splicing kinetics are identical for Genes 1 and 2. Thus, splicing for Gene 2 is represented by:

$$T_2 \xrightarrow{\sigma} M_2 + x\ X_2$$

*Translation*

Translation of the mature mRNA transcript for Gene 1 is represented by the following reaction:

$$M_1 \xrightarrow{\kappa} M_1 + A_1 + R_1$$

where $\kappa$ represents the translation rate constant. We assume the translation kinetics are identical for Genes 1 and 2. Thus, translation for Gene 2 is represented by:

$$M_2 \xrightarrow{\kappa} M_2 + A_2 + R_2$$

*Degradation*

Protein degradation is modeled simply as first-order decay with a rate constant, $\gamma$. Degradation of mRNA species is dependent on microRNA level, and is modeled with the following reactions:

$$M_1 \xrightarrow{\delta} \varnothing$$

$$X_2 + M_1 \underset{k_{X2,unbind}}{\overset{k_{X2,bind}}{\longleftrightarrow}} C_{X2,M1} \xrightarrow{k_{2,decay}} X_2$$

where $\delta$ is the basal rate constant for mRNA decay, $k_{2,decay}$ is the rate constant for microRNA-mediated decay, $k_{X2,bind}$ is the rate constant for microRNA association, and $k_{X2,unbind}$ is the rate constant for microRNA dissociation. The rate of degradation of $M_1$ is:

$$f_{1,decay} = \delta M_1 + k_{X2,bind} X_2 M_1 - k_{X2,unbind} C_{X2,M1}$$

Assuming that the dynamics of microRNA binding are fast compared to degradation, we can solve for the concentration of the complex:

$$C_{X2,M_1} = \frac{k_{X2,bind} X_2 M_1}{k_{X2,unbind} + k_{2,decay}} = \frac{X_2 M_1}{K_{M,2}}$$

where $K_{M,2}$ is the Michaelis constant for microRNA-mediated degradation. The rate of $M_1$ decay can then be simplified to a function of $M_1$ and $X_2$:

$$f_{1,decay}(M_1, X_2) = \Big(\delta + \xi_2 X_2\Big) M_1$$

where $\xi_2 = k_{2,decay}/K_{M,2}$ and represents the microRNA repression strength. Similarly, the rate of $M_2$ decay is represented as:

$$f_{2,decay}(M_2, X_1) = \Big(\delta + \xi_1 X_1\Big) M_2$$

If we consider that unspliced transcripts may also be knocked down by microRNA expression, we obtain the following rates of $T_1$ and $T_2$ decay:

$$f_{1,decay}(T_1, X_2) = \Big(\delta + \xi_2 X_2\Big) T_1$$

$$f_{2,decay}(T_2, X_1) = \Big(\delta + \xi_1 X_1\Big) T_2$$

*Full deterministic model*

Incorporating the simplifications above, we obtain the following differential equations to describe the two gene system:

$$\frac{dT_1}{dt} = G_{1,tot} \frac{\alpha_1 + \frac{\beta_1 A_1}{K_{D,A1}}}{1 + \frac{A_1}{K_{D,A1}} + \frac{R_2}{K_{D,R2}}} - \Big(\sigma + \delta + \xi_2 X_2\Big) T_1 \tag{S1}$$

$$\frac{dT_2}{dt} = G_{2,tot} \frac{\alpha_2 + \frac{\beta_2 A_2}{K_{D,A2}}}{1 + \frac{A_2}{K_{D,A2}} + \frac{R_1}{K_{D,R1}}} - \Big(\sigma + \delta + \xi_1 X_1\Big) T_2 \tag{S2}$$

$$\frac{dM_1}{dt} = \sigma T_1 - \Big(\delta + \xi_2 X_2\Big) M_1 \tag{S3}$$

$$\frac{dM_2}{dt} = \sigma T_2 - \Big(\delta + \xi_1 X_1\Big) M_2 \tag{S4}$$

$$\frac{dX_1}{dt} = x\sigma T_1 - \delta X_1 \tag{S5}$$

$$\frac{dX_2}{dt} = x\sigma T_2 - \delta X_2 \tag{S6}$$

$$\frac{dA_1}{dt} = \kappa M_1 - \gamma A_1 \tag{S7}$$

$$\frac{dA_2}{dt} = \kappa M_2 - \gamma A_2 \tag{S8}$$

$$\frac{dR_1}{dt} = \kappa M_1 - \gamma R_1 \tag{S9}$$

$$\frac{dR_2}{dt} = \kappa M_2 - \gamma R_2 \tag{S10}$$

where $x = 1$ for circuits employing microRNA-based repression. This variable is otherwise equal to zero. Base parameter values are listed in Table S1.

#### Modeling of shared transcriptional co-activators

To account for limited transcriptional co-activator resources, the differential equations governing the primary RNA transcript dynamics are updated to:

$$\frac{dT_1}{dt} = G_{1,tot}\frac{1}{1+\frac{A_1}{K_{M1}}+c\frac{A_2}{K_{M2}}}\frac{\alpha_1+\frac{\beta_1 A_1}{K_{D,A1}}}{1+\frac{A_1}{K_{D,A1}}+\frac{R_2}{K_{D,R2}}}-\left(\sigma+\delta+\xi_2 X_2\right)T_1 \tag{S11}$$

$$\frac{dT_2}{dt} = G_{2,tot}\frac{1}{1+c\frac{A_1}{K_{M1}}+\frac{A_2}{K_{M2}}}\frac{\alpha_2+\frac{\beta_2 A_2}{K_{D,A2}}}{1+\frac{A_2}{K_{D,A2}}+\frac{R_1}{K_{D,R1}}}-\left(\sigma+\delta+\xi_1 X_1\right)T_2, \tag{S12}$$

where $K_{M1}$ and $K_{M2}$ represent the co-activator dissociation constants for each of the transcriptional activators $A_1$ and $A_2$, respectively. For derivation of the additional multiplicative terms in Equations (S11) and (S12) compared to Equations (S1) and (S2), see Jones and Qian *et al* [31]. For circuits with identical activation domains, let $c = 1$ to represent a shared resource pool. For circuits with different activation domains, let $c = 0$ to represent separate resource pools.

We wish to use the models above to observe how having non-identical activation domains (that is, when $c = 0$) can be a detriment to bistable circuit design. To this end, we consider the case $c = 0$, and observe how the number of stable steady states is affected by gene dosage (Fig. S1). As can be seen, both identical and non-identical activation domains start with a single steady state; after a threshold gene dosage, bistability (two stable steady states, with one unstable steady state for a total of three) emerges in both systems. With identical domains (right), this bistability is maintained *ad infinitum* as gene dosages are increased in coordination with one another. However, with non-identical domains, there is a loss in bistability as gene dosages increases. After a threshold gene dosage, the non-identical domain system is entirely monostable.

In the model, one can think of the new multiplicative term as a scaling factor for the old bistable system model. When $c = 1$, these scaling factors change together; when $c = 0$, they shift independently of one another and make the balance of factors needed to achieve bistability in the model untenable.

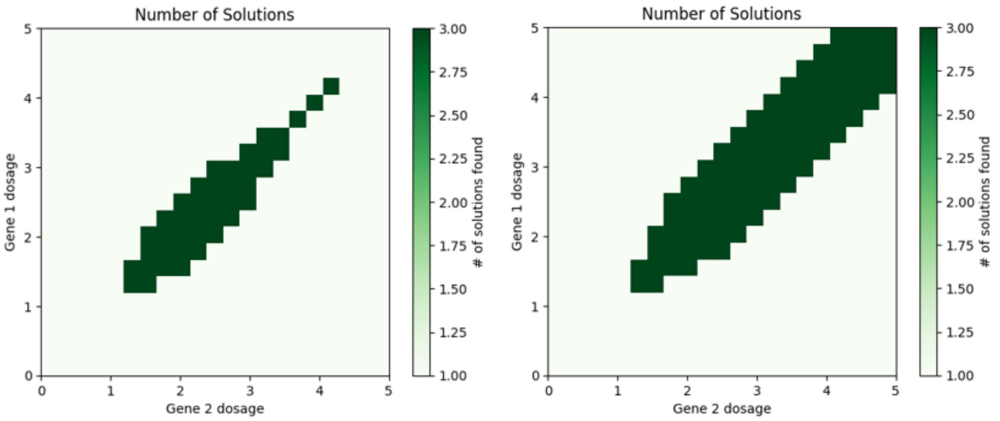


**Figure S1.** Plots of the number of steady states in the system defined by (S11) and (S12) when $c = 0$ (left) or $c = 1$ (right). Gene dosages are shown in a log scale. Parameters used to generate this figure are: $Gi,tot = \sigma+\delta+\xi_i X_i = \alpha_i = 1$, $\frac{\beta_i}{K_{D,Ai}} = 100$, $K_{D,Ai} = 1000$, $K_{D,Ri} = 100$, $K_{Mi} = 10000$.

#### Supplementary Figures

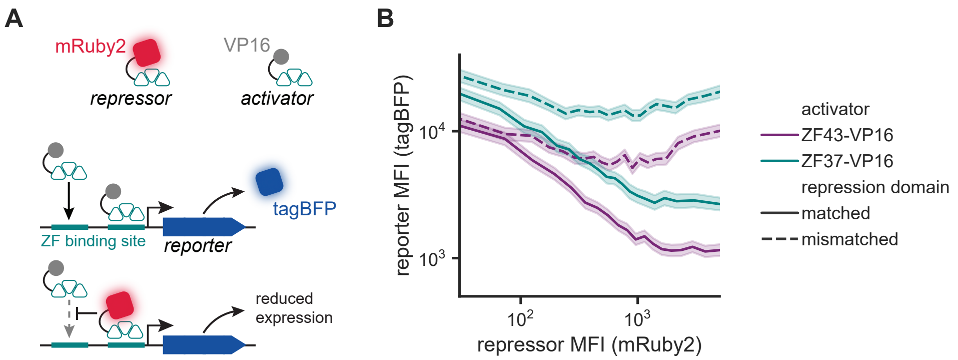


**Figure S2. ZF-based repression tunes reporter expression in a dose-dependent manner.**
A. The ZF-based repressors used here consist of a ZF DBD fused to a fluorescent protein. Upon binding to its cognate promoter, the ZF-based repressor prevents adjacent binding of activators through steric hindrance.
B. Reporter expression (tagBFP MFI) as a function of repressor expression level (mRuby2 MFI) for one representative biological replicate. Constructs were poly-transfected into HEK293T cells and analyzed via flow cytometry at 2 dpt. Data was separated into 20 equal-quantile bins based on repressor expression. Lines represent the geometric mean of the MFI for each bin. Shaded regions represent the 95% confidence interval. All data is in arbitrary units from a flow cytometer.

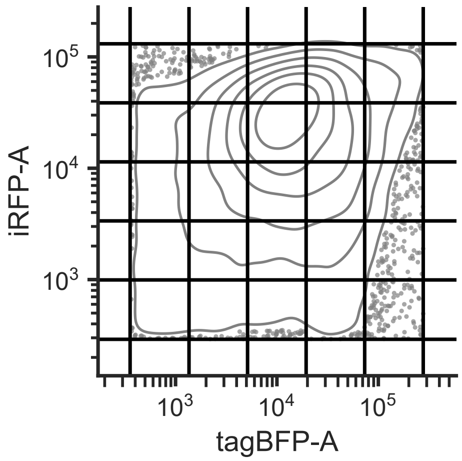


**Figure S3. Bins used in poly-transfection characterization of circuit bias.** Density plot of marker gene expression (tagBFP and iRFP) for HEK293T cells poly-transfected with the circuit depicted in Figure 2A. Points represent the 5% of probability mass lying outside the lowest iso-proportion contour. Data is shown for one representative biological replicate. Black lines indicate equal width copy number bins used in further poly-transfection analysis of the circuits. All data is in arbitrary units from a flow cytometer.

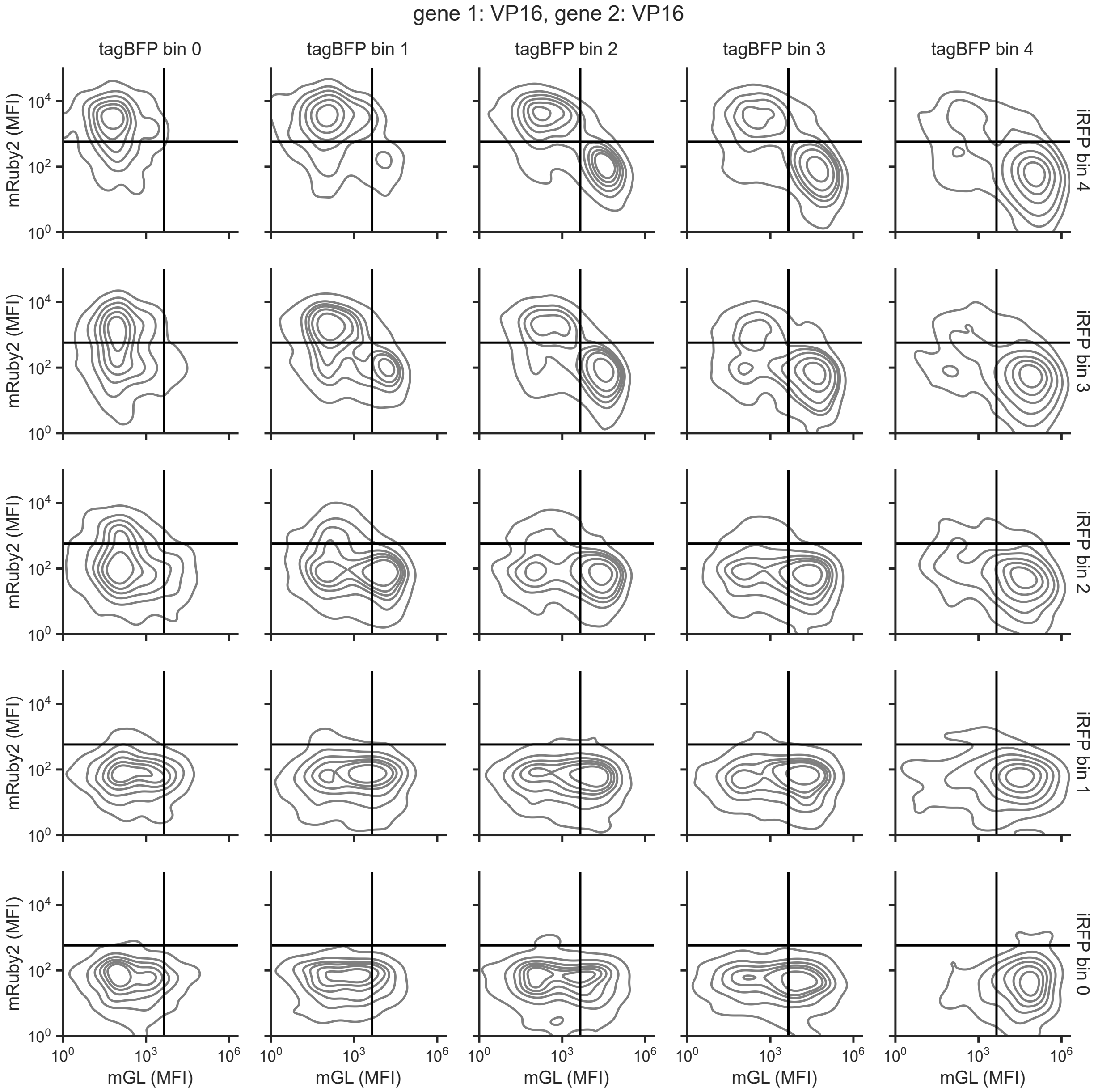


**Figure S4. Poly-transfection characterization of the initial circuit design.** Density plots of circuit gene expression within varying copy number bins for HEK293T cells poly-transfected with the circuit depicted in Figure 2A. Data is shown for one representative biological replicate. Black lines indicate gates used in calculating the bias scores reported in Figure 3. All data is in arbitrary units from a flow cytometer.

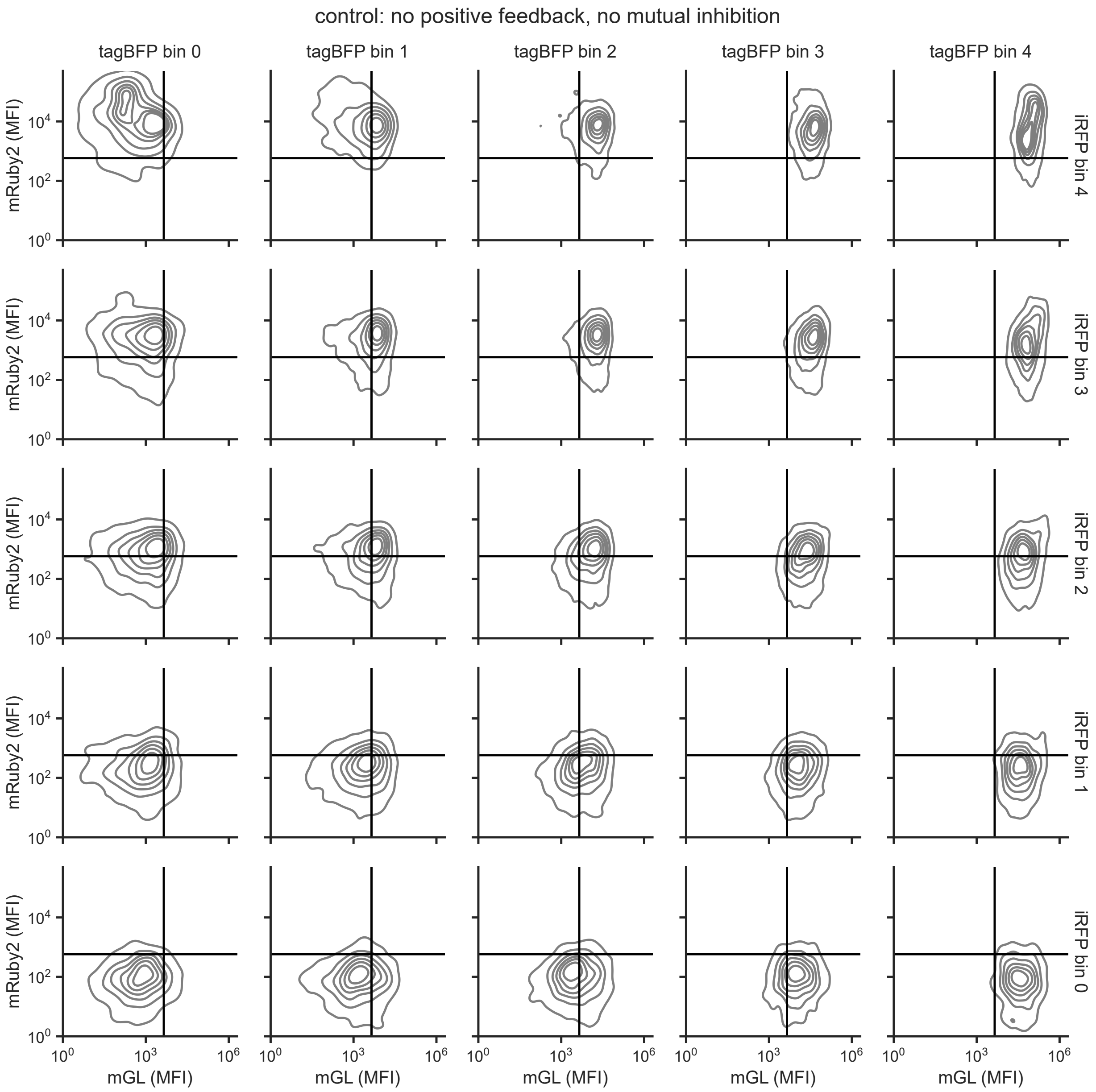


**Figure S5. Poly-transfection characterization of the control condition.** Density plots of circuit gene expression within varying copy number bins for HEK293T cells poly-transfected with the control circuit depicted in Figure 2. Data is shown for one representative biological replicate. All data is in arbitrary units from a flow cytometer.

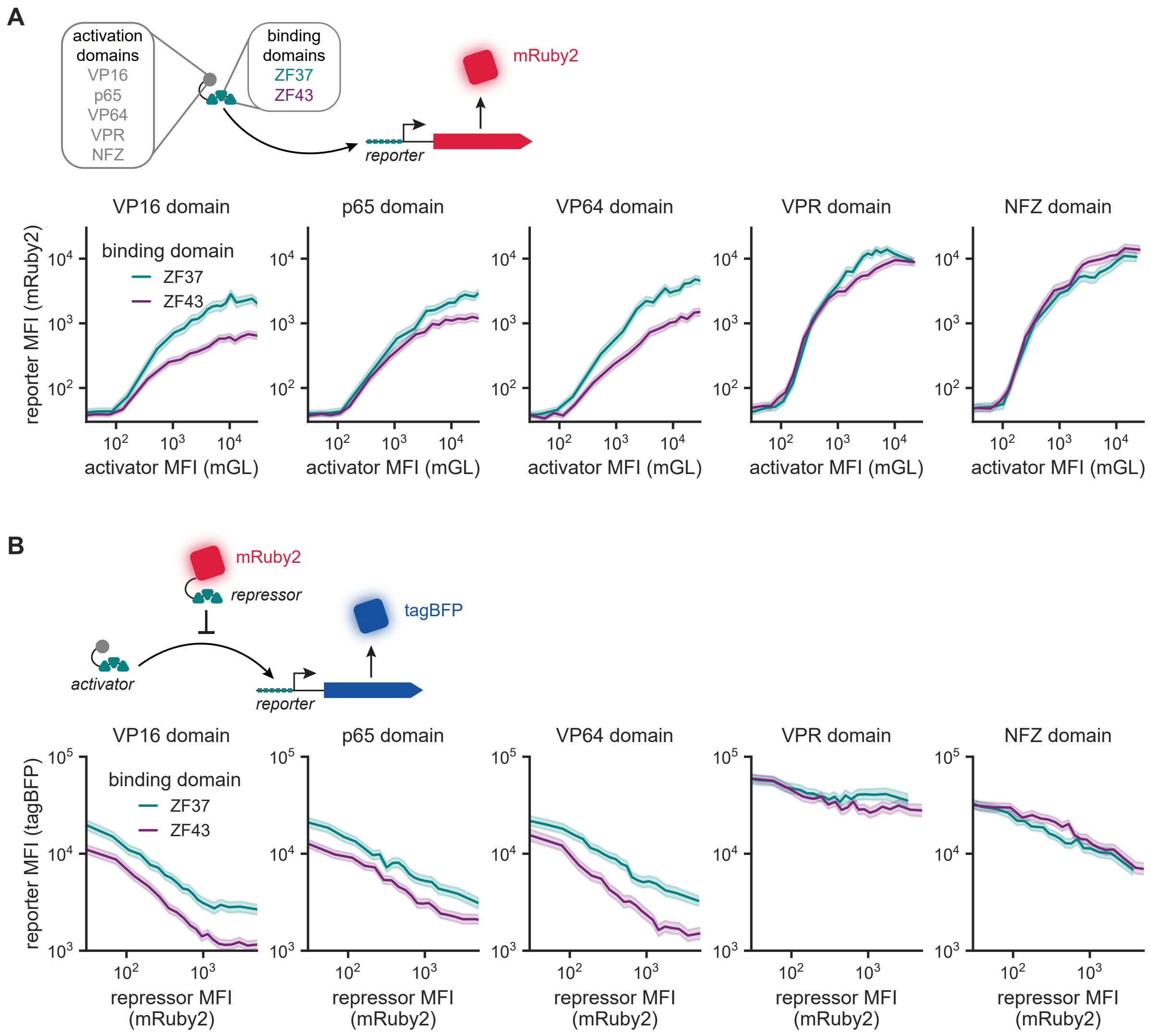


**Figure S6. Characterization of ZF-based activation and repression for varying activation domains.** (A) Reporter expression (mRuby2 MFI) as a function of activator expression level (mGL MFI) for ZF-based activators containing varying activation domains. Data shown is one representative biological replicate. Constructs were poly-transfected into HEK293T cells and analyzed via flow cytometry at 2 dpt. Data was separated into 20 equal-quantile bins based on activator expression. (B) Reporter expression (tagBFP MFI) as a function of repressor expression level (mRuby2 MFI) for ZF-based repressors paired with varying activation domains. Data shown is one representative biological replicate. Constructs were poly-transfected into HEK293T cells and analyzed via flow cytometry at 2 dpt. Data was separated into 20 equal-quantile bins based on repressor expression. Lines represent the geometric mean of the MFI for each bin. Shaded regions represent the 95% confidence interval. All data is in arbitrary units from a flow cytometer.

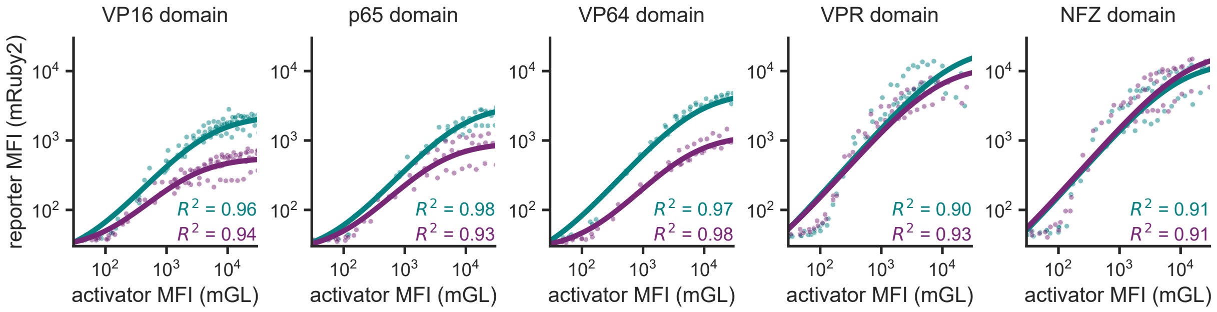


**Figure S7. Fitting of activator poly-transfection data.** Binned data from Figure S6 was fit the following equation: $\mathrm{MFI}_{reporter} = \alpha_0 + \frac{\alpha \cdot \mathrm{MFI}_{activator}}{K + \mathrm{MFI}_{activator}}$, where $\mathrm{MFI}_{reporter}$ is the measured reporter fluorescence, $\mathrm{MFI}_{activator}$ is the measured activator fluorescence, $\alpha_0$ represents leaky reporter expression, $\alpha$ represents maximum activated reporter expression, and $K$ represents the activator expression level at which the half maximal reporter expression is achieved. Points represent geometric means of individual activator bins taken from at least three biological replicates. Solid line represents the fitted model.

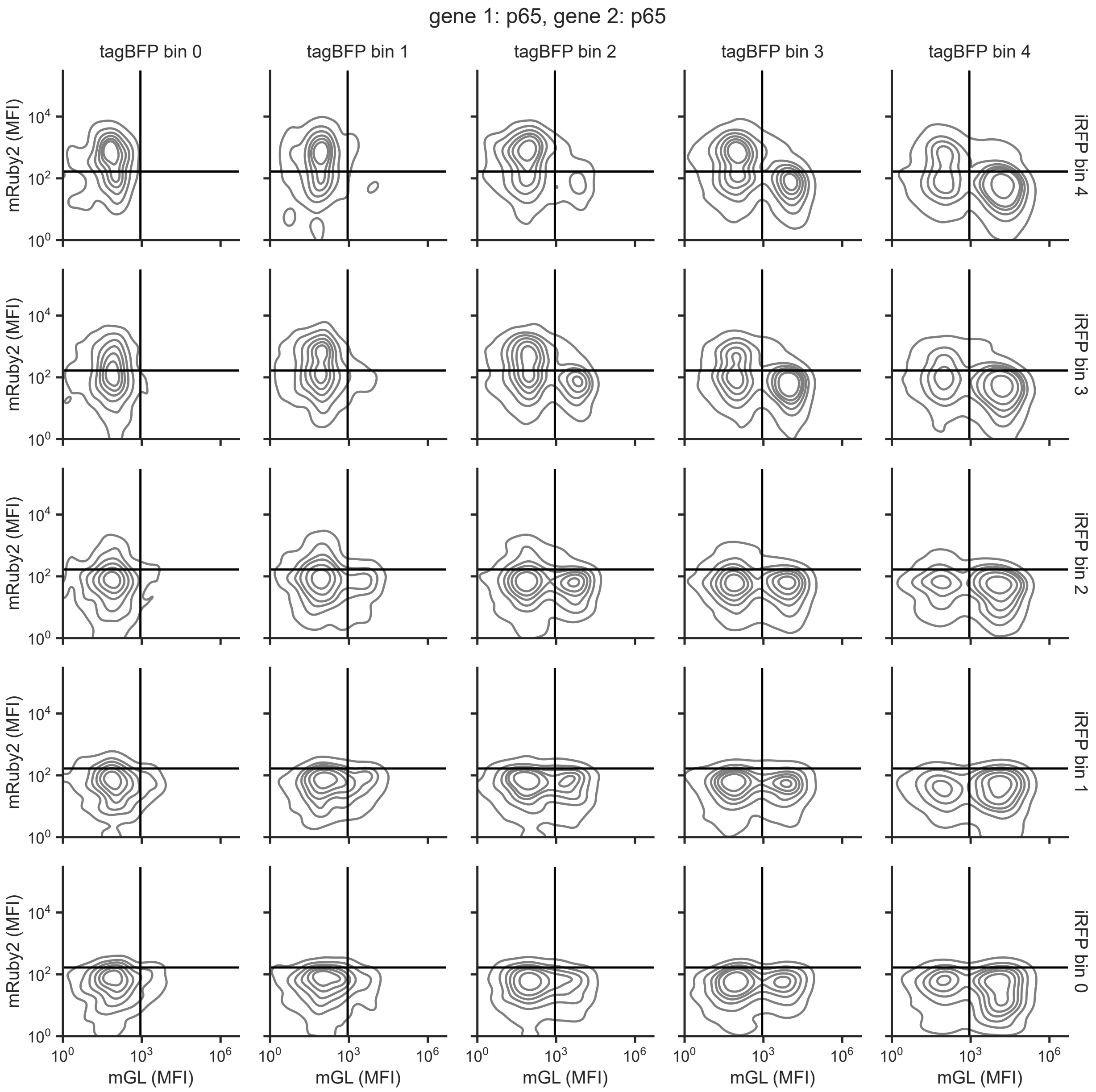


**Figure S8. Poly-transfection characterization of the circuit with two p65 activation domains.** Density plots of circuit gene expression within varying copy number bins for HEK293T cells poly-transfected with the circuit containing two p65-based activators. Data is shown for one representative biological replicate. Black lines indicate gates used in calculating the bias scores reported in Figure 4C. All data is in arbitrary units from a flow cytometer.

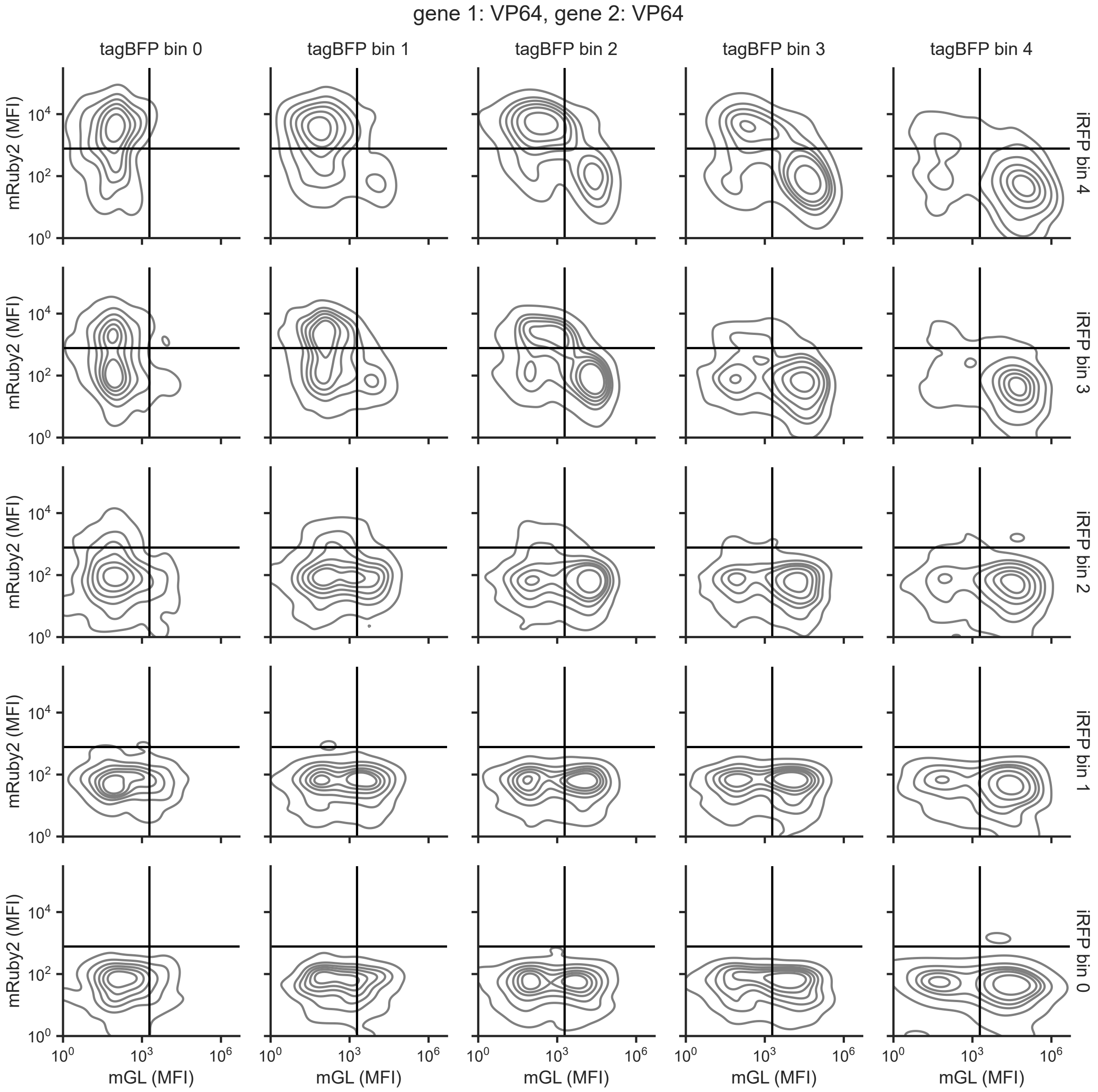


**Figure S9. Poly-transfection characterization of the circuit with two VP64 activation domains.** Density plots of circuit gene expression within varying copy number bins for HEK293T cells poly-transfected with the circuit containing two VP64-based activators. Data is shown for one representative biological replicate. Black lines indicate gates used in calculating the bias scores reported in Figure 4C. All data is in arbitrary units from a flow cytometer.

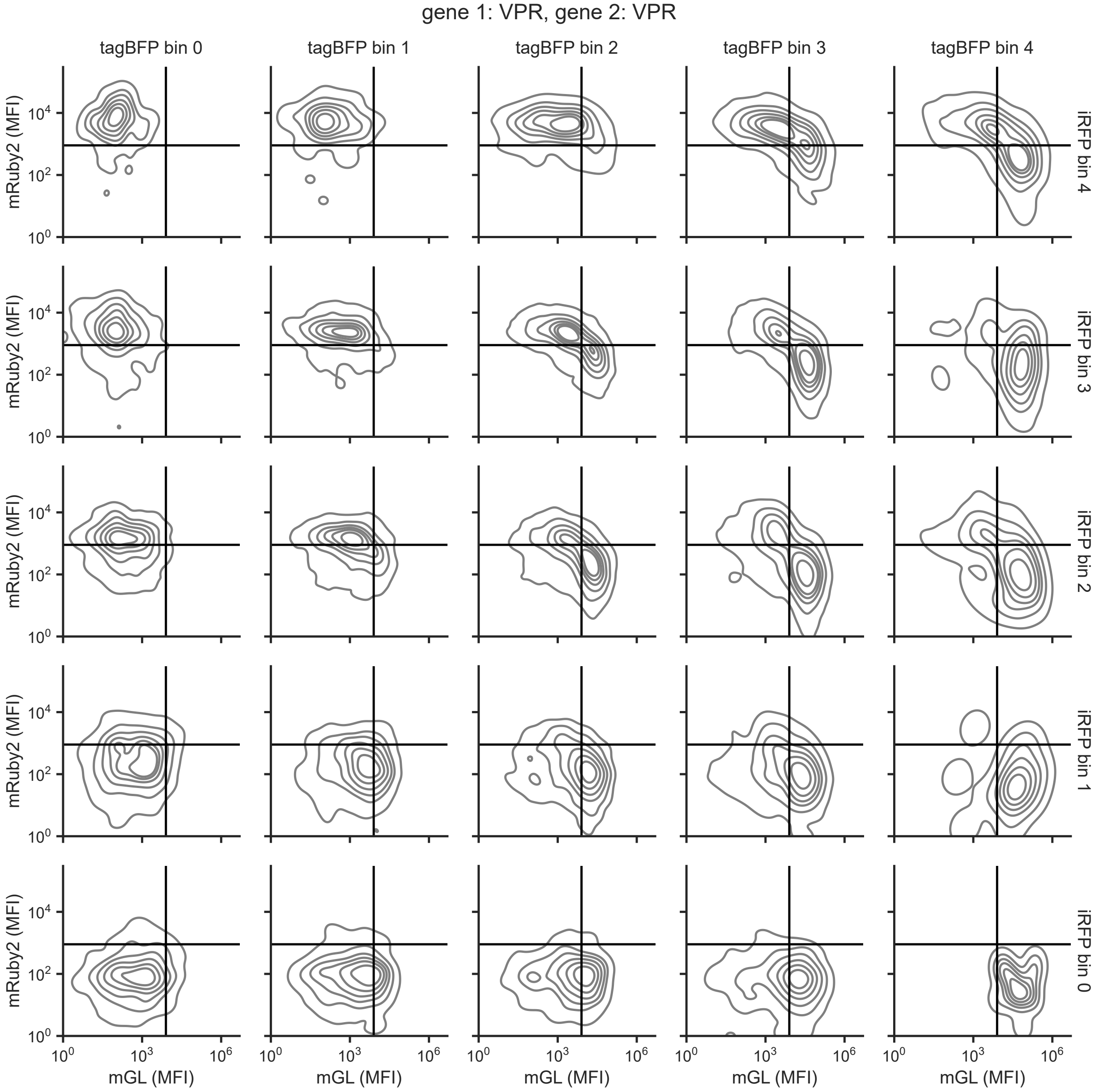


**Figure S10. Poly-transfection characterization of the circuit with two VPR activation domains.** Density plots of circuit gene expression within varying copy number bins for HEK293T cells poly-transfected with the circuit containing two VPR-based activators. Data is shown for one representative biological replicate. Black lines indicate gates used in calculating the bias scores reported in Figure 4C. All data is in arbitrary units from a flow cytometer.

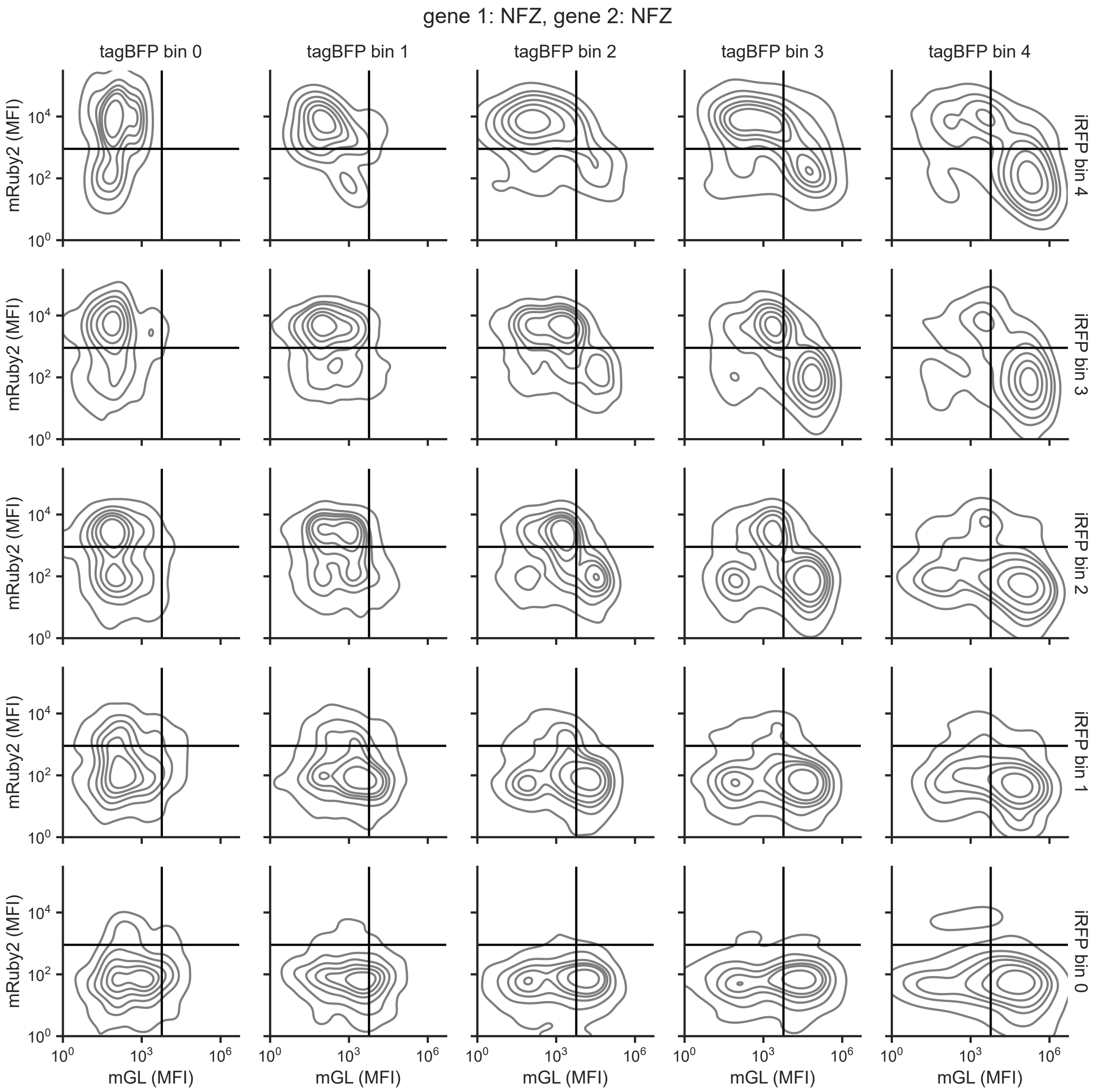


**Figure S11. Poly-transfection characterization of the circuit with two NFZ activation domains.** Density plots of circuit gene expression within varying copy number bins for HEK293T cells poly-transfected with the circuit containing two NFZ-based activators. Data is shown for one representative biological replicate. Black lines indicate gates used in calculating the bias scores reported in Figure 4C. All data is in arbitrary units from a flow cytometer.

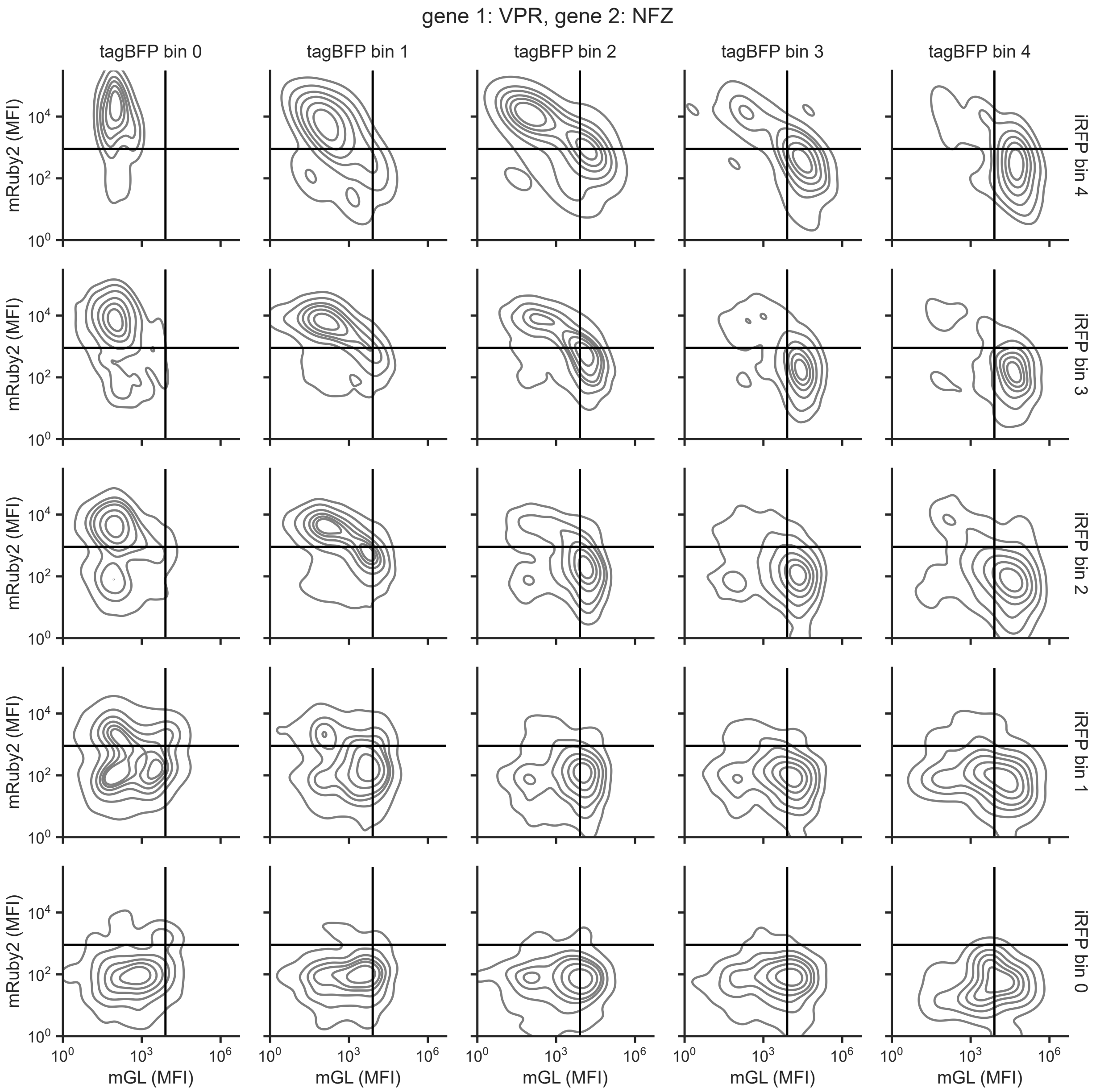


**Figure S12. Poly-transfection characterization of the circuit with a VPR domain on gene 1 and an NFZ domain on gene 2.** Density plots of circuit gene expression within varying copy number bins for HEK293T cells poly-transfected with the circuit containing VPR-ZF37 and NFZ-ZF43. Data is shown for one representative biological replicate. All data is in arbitrary units from a flow cytometer.

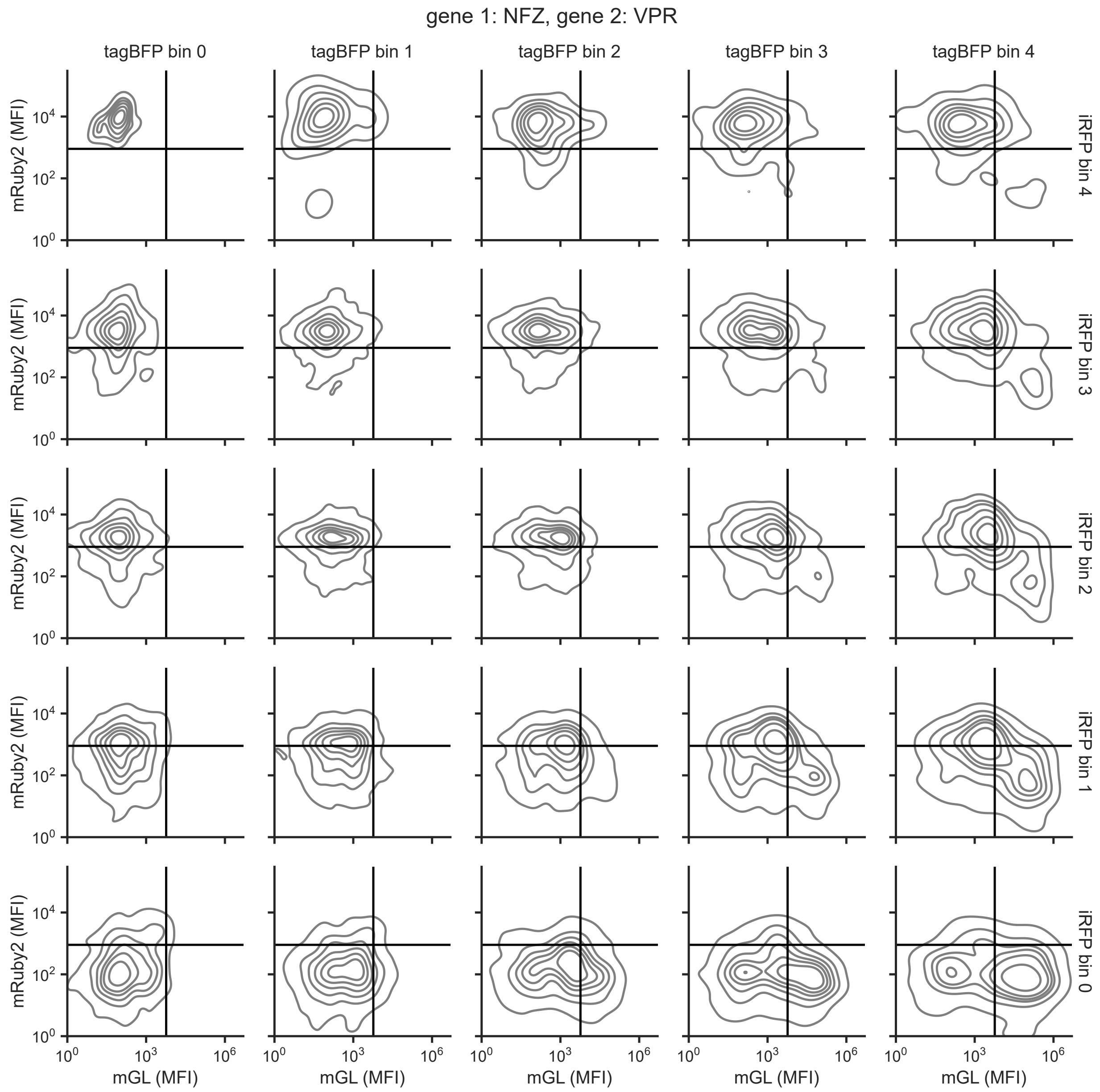


**Figure S13. Poly-transfection characterization of the circuit with an NFZ domain on gene 1 and a VPR domain on gene 2.** Density plots of circuit gene expression within varying copy number bins for HEK293T cells poly-transfected with the circuit containing NFZ-ZF37 and VPR-ZF43. Data is shown for one representative biological replicate. All data is in arbitrary units from a flow cytometer.

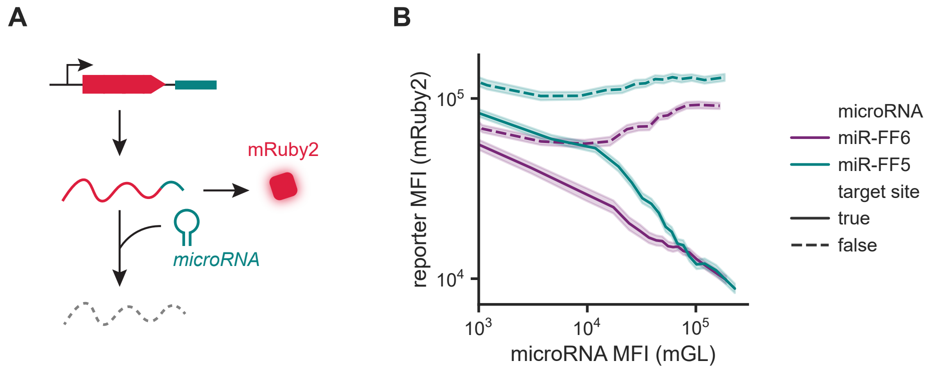


**Figure S14. Synthetic microRNAs reduce target expression in a dose-dependent manner.**
A. The synthetic microRNA sequences tested here include miR-FF5 and miR-FF6. Each was paired with reporters with and without a cognate target site in the 3' UTR and characterized in HEK293T cells via poly-transfection.
B. Reporter expression (mRuby2 MFI) as a function of microRNA expression level (mGL MFI) for microRNAs encoded within a synthetic intron containing the miR-30 backbone. Data is shown for one representative biological replicate. Constructs were poly-transfected into HEK293T cells and analyzed via flow cytometry at 2 dpt. Data was separated into 20 equal-quantile bins based on microRNA expression. Lines represent the geometric mean of the MFI for each bin. Shaded regions represent the 95% confidence interval. All data is in arbitrary units from a flow cytometer.

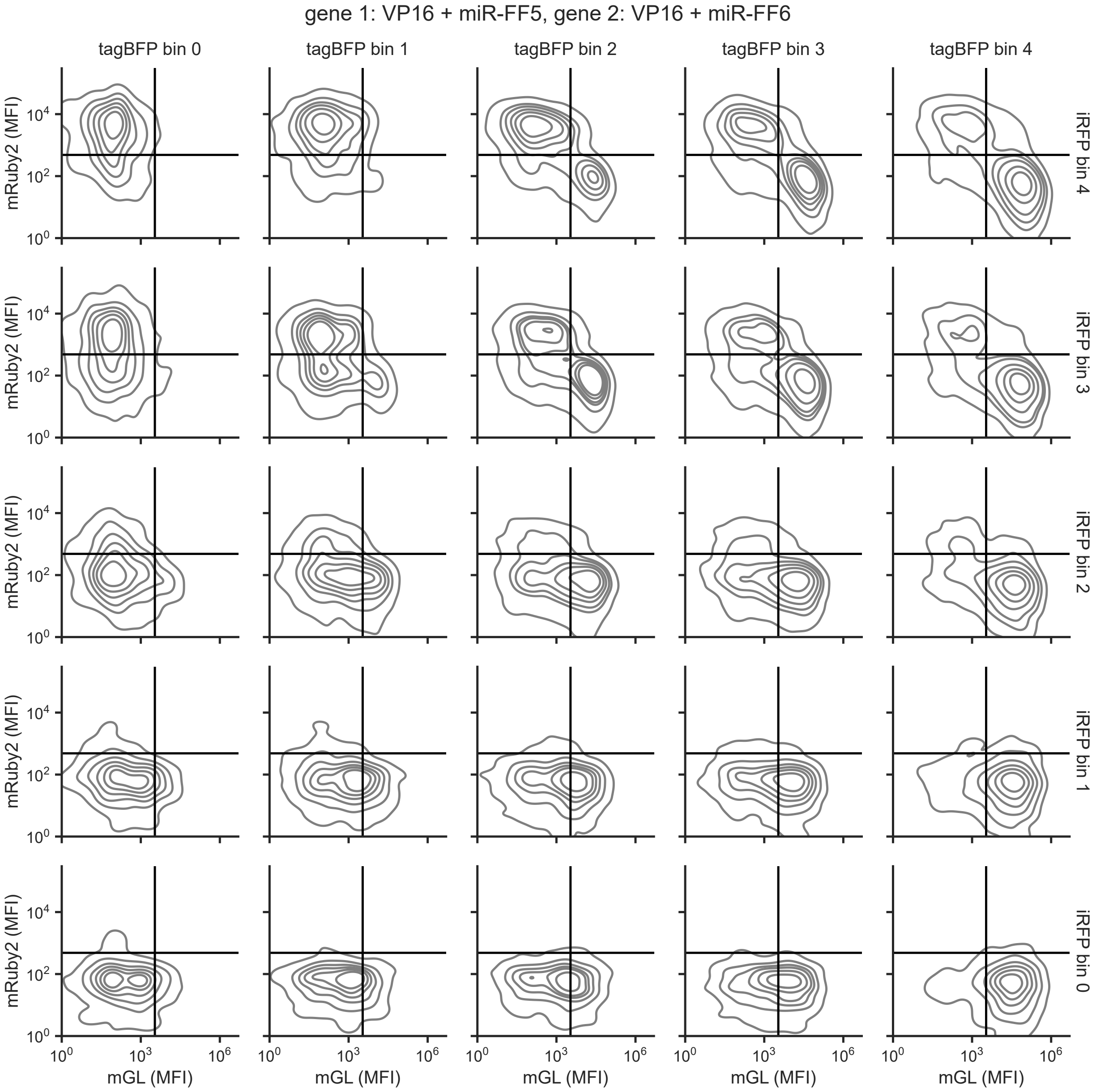


**Figure S15. Poly-transfection characterization of circuit with VP16-based activation and additional microRNA-based mutual repression.** Density plots of circuit gene expression within varying copy number bins for HEK293T cells poly-transfected with the VP16 circuit depicted in Figure 6C. Data is shown for one representative biological replicate. Black lines indicate gates used in calculating the bias scores reported in Figure 6D. All data is in arbitrary units from a flow cytometer.

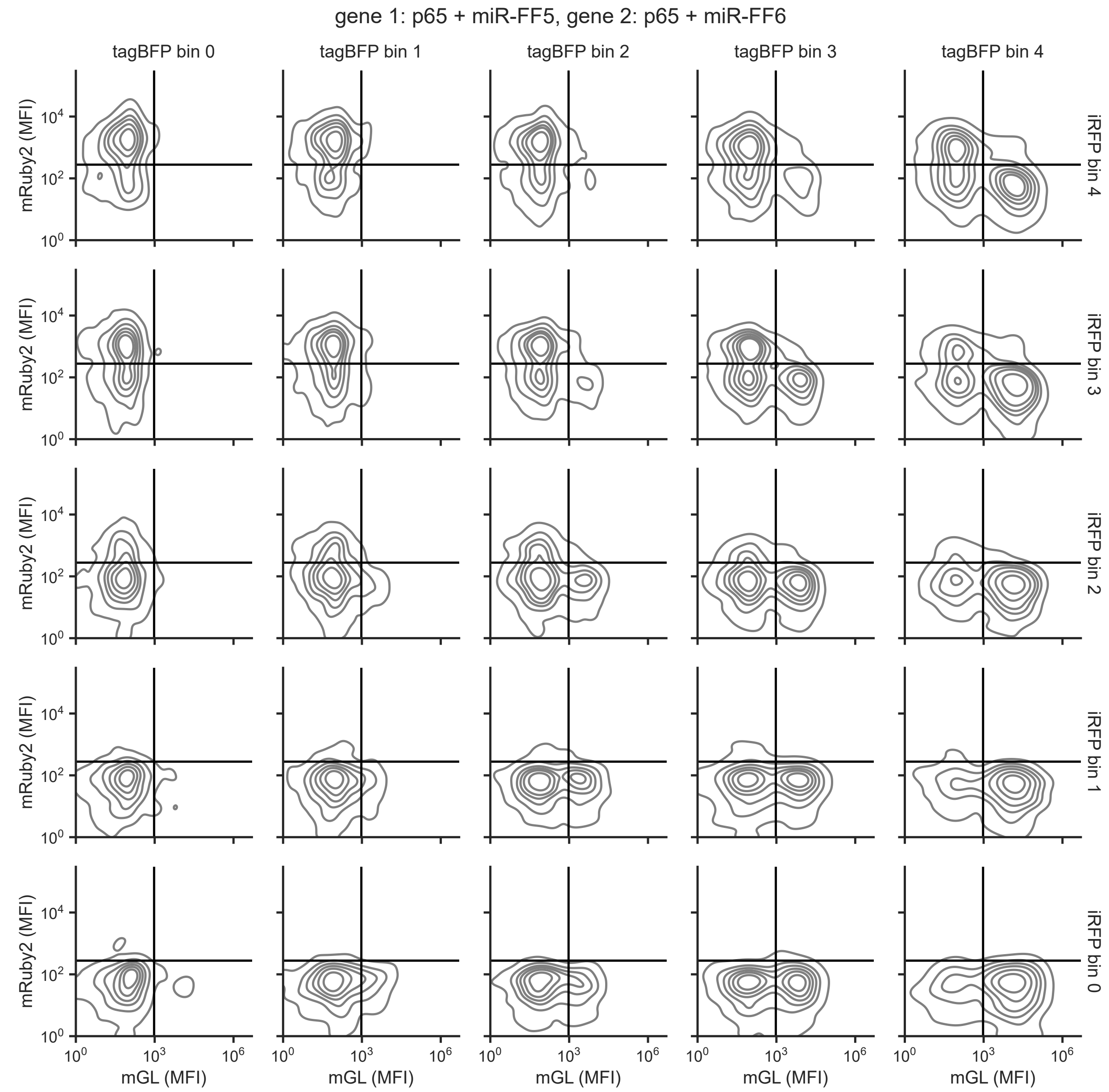


**Figure S16. Poly-transfection characterization of circuit with p65-based activation and additional microRNA-based mutual repression.** Density plots of circuit gene expression within varying copy number bins for HEK293T cells poly-transfected with the p65 circuit depicted in Figure 6C. Data is shown for one representative biological replicate. Black lines indicate gates used in calculating the bias scores reported in Figure 6D. All data is in arbitrary units from a flow cytometer.

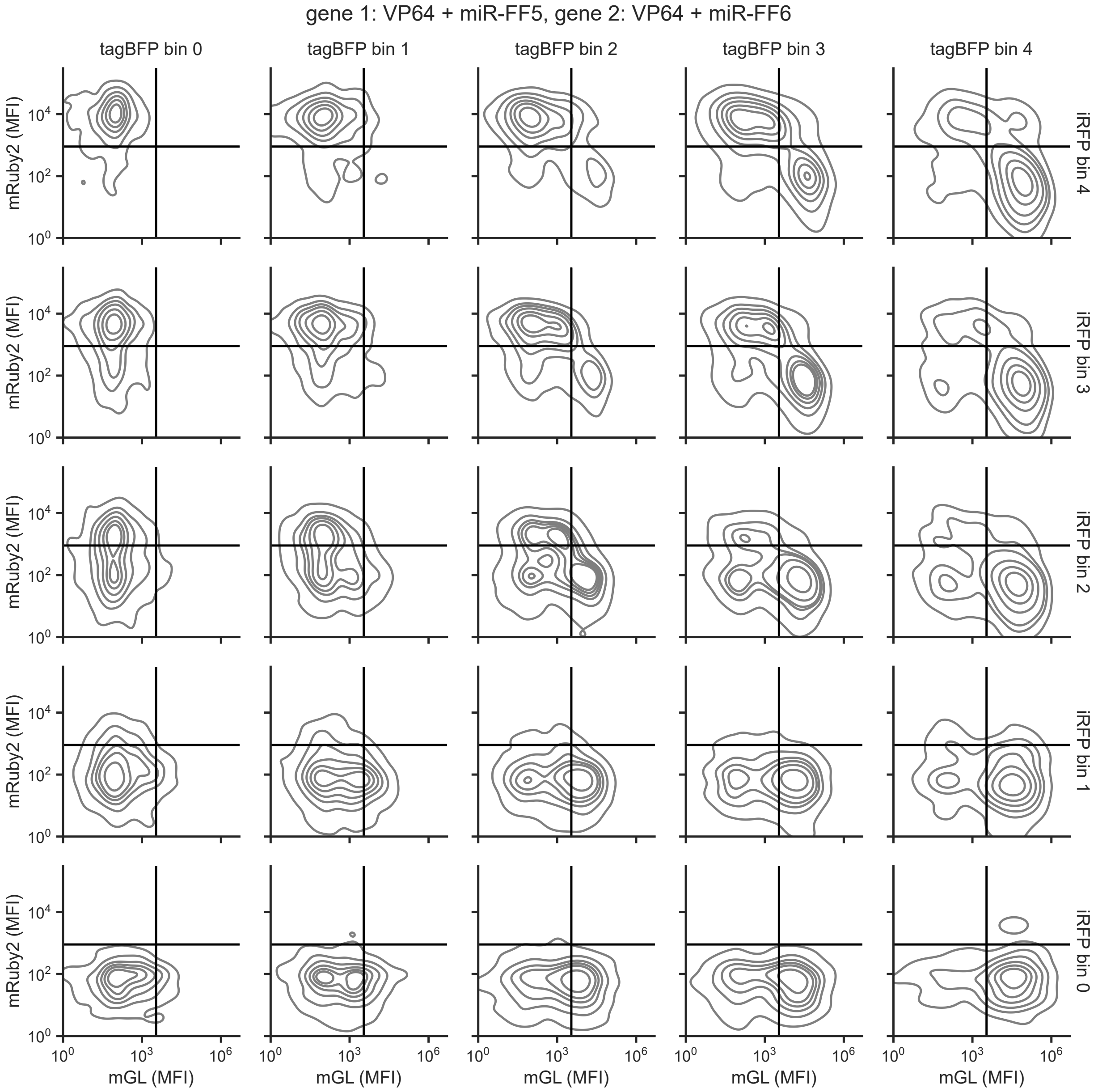


**Figure S17. Poly-transfection characterization of circuit with VP64-based activation and additional microRNA-based mutual repression.** Density plots of circuit gene expression within varying copy number bins for HEK293T cells poly-transfected with the VP64 circuit depicted in Figure 6C. Data is shown for one representative biological replicate. Black lines indicate gates used in calculating the bias scores reported in Figure 6D. All data is in arbitrary units from a flow cytometer.

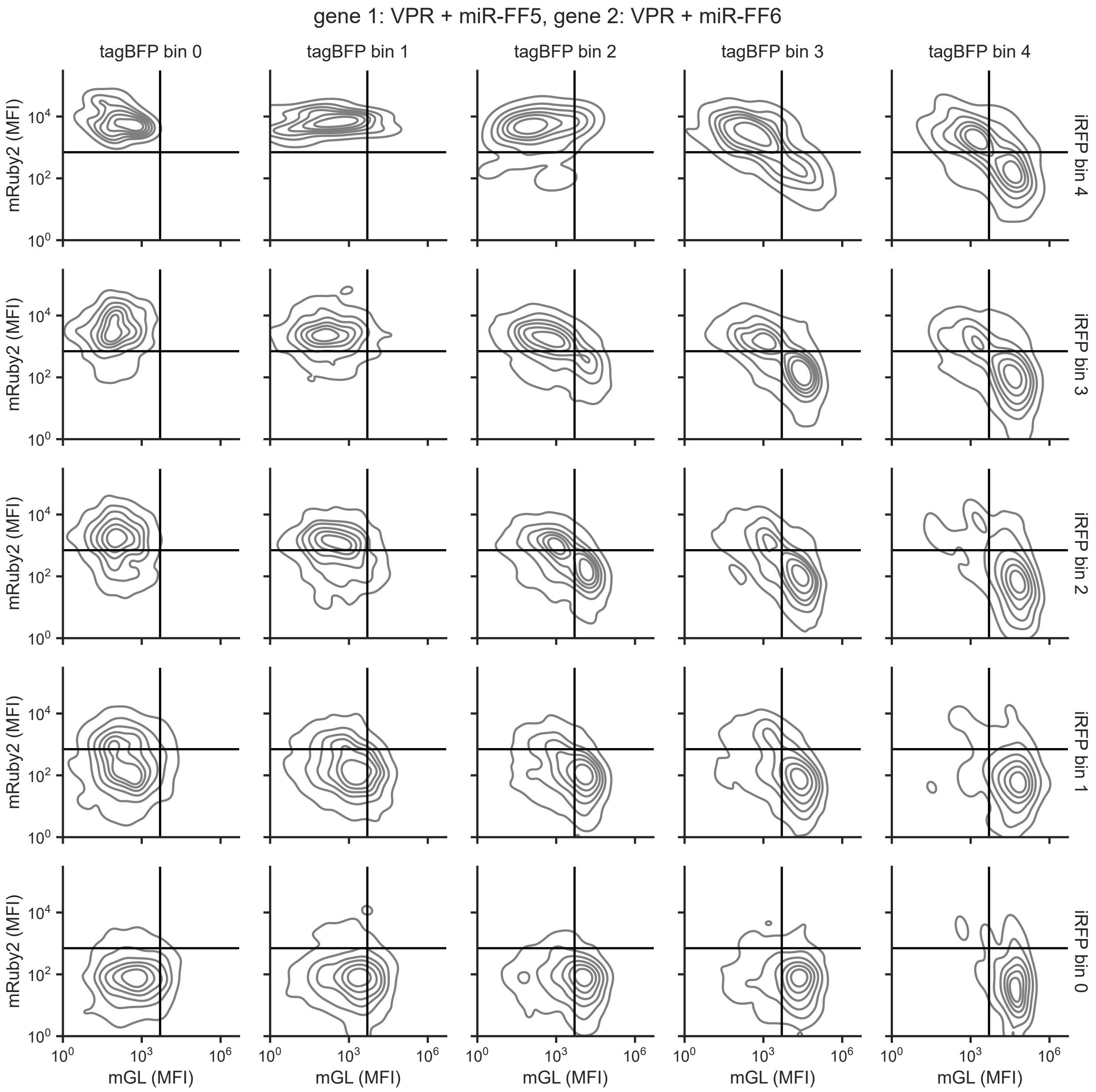


**Figure S18. Poly-transfection characterization of circuit with VPR-based activation and additional microRNA-based mutual repression.** Density plots of circuit gene expression within varying copy number bins for HEK293T cells poly-transfected with the VPR circuit depicted in Figure 6C. Data is shown for one representative biological replicate. Black lines indicate gates used in calculating the bias scores reported in Figure 6D. All data is in arbitrary units from a flow cytometer.

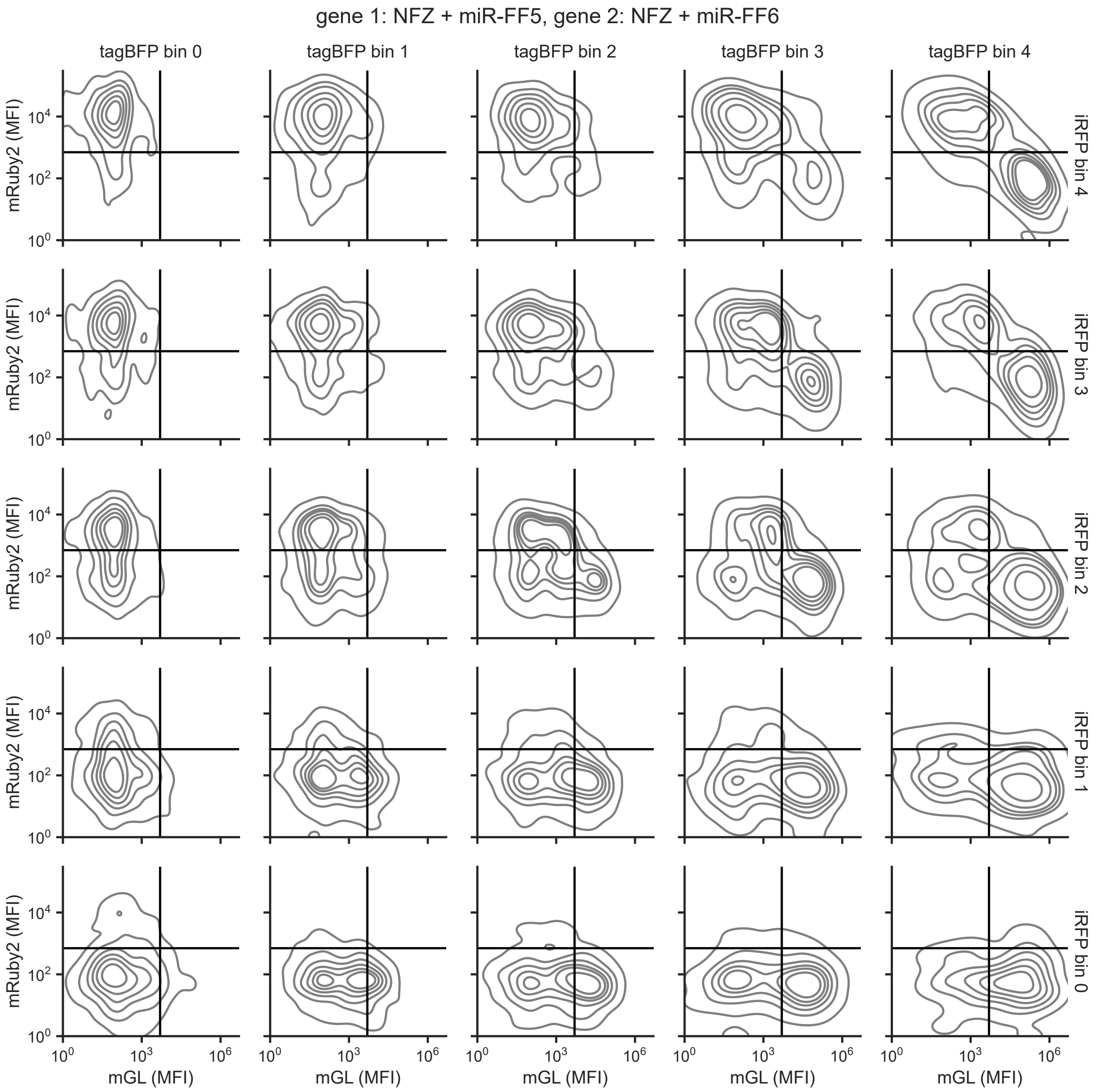


**Figure S19. Poly-transfection characterization of circuit with NFZ-based activation and additional microRNA-based mutual repression.** Density plots of circuit gene expression within varying copy number bins for HEK293T cells poly-transfected with the NFZ circuit depicted in Figure 6C. Data is shown for one representative biological replicate. Black lines indicate gates used in calculating the bias scores reported in Figure 6D. All data is in arbitrary units from a flow cytometer.

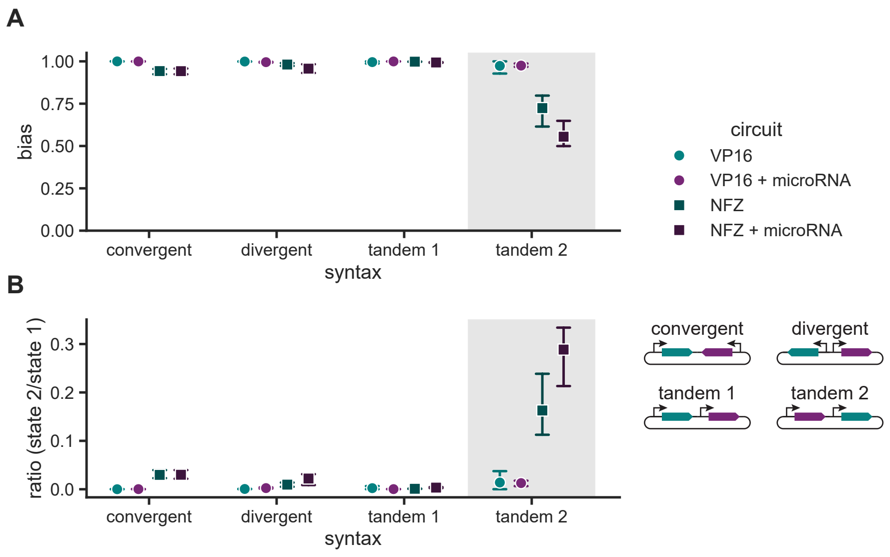


**Figure S20. Additional characterization data for single-cargo DAC designs.** (**A**) Circuit bias for all single-cargo DAC designs, calculated as the difference between the numbers of mGL+ and mRuby2+ cells divided by their sum. (**B**) Ratio of cells in state 2 (mRuby2+) to cells in state 1 (mGL+) for all single-cargo DAC designs. Data points represent the mean of three biological replicates, and error bars represent the 95% confidence interval. Shaded regions indicate the DAC designs displayed in Figures 7E and 7F.

### Supplementary Tables

**Table S1. Base parameter values used in the deterministic circuit model.** Concentrations are estimated assuming a cell diameter of 14 $\mu$M and a spherical volume of approximately 1.5 pL [45].

| Parameter | Description | Value | Units | Source |
|---|---|---|---|---|
| $G_{1,tot}$ and $G_{2,tot}$ | Gene copy number | $10^{-2}$ to $10^{5}$, 1 (default) | pM | [46, 47] |
| $\alpha_1$ and $\alpha_2$ | Basal transcription | $2 \cdot 10^{-1}$ | 1 / s | [13] |
| $\beta_1$ and $\beta_2$ | Activated transcription | 6 | 1 / s | [13] |
| $K_{D,A1}$ and $K_{D,A2}$ | Activator dissociation | 50 | pM | [25] |
| $K_{D,R1}$ and $K_{D,R2}$ | Repressor dissociation | 20 | pM | [12] |
| $\xi_1$ and $\xi_2$ | microRNA repression | $4 \cdot 10^{-8}$ | 1 / pM · s | [48, 49] |
| $\sigma$ | Splicing | $2 \cdot 10^{-3}$ | 1 / s | [50] |
| $\kappa$ | Translation | $3 \cdot 10^{-4}$ | 1 / s | [48, 51] |
| $\delta$ | RNA decay | $3 \cdot 10^{-4}$ | 1 / s | [51] |
| $\gamma$ | Protein decay | $1 \cdot 10^{-4}$ | 1 / s | [51] |
| $K_M$ | Co-activator dissociation | $1 \cdot 10^{5}$ | pM | [52] |

**Table S2. Fitted parameters for each transcriptional activator design and the corresponding parameters used in deterministic modeling.** Experimental flow-cytometry data (Figures S6 and S7) was fit to the following model: $\text{MFI}_{reporter} = \alpha_0 + \frac{\alpha \cdot \text{MFI}_{activator}}{K + \text{MFI}_{activator}}$, where $\text{MFI}_{reporter}$ is the measured reporter fluorescence, $\text{MFI}_{activator}$ is the measured activator fluorescence, $\alpha_0$ represents leaky reporter expression, $\alpha$ represents maximum activated reporter expression, and $K$ represents the activator expression level at which the half maximal reporter expression is achieved. Reporter data was log-transformed for fitting to reduce bias of the least-squares fit towards high numbers. Values reported here are the fitted parameters ± the standard deviation. To model varying activation domains, the $\beta_i$ parameters (Table S1) were scaled based on the fitted $\alpha$ parameter for VP16-ZF37.

| Binding domain: ZF37 | | | | |
|---|---|---|---|---|
| Activation domain | $\alpha_0$ (a.u.) | $\alpha$ (a.u.) | $K$ (a.u.) | $\beta_1$ (1 / s) |
| VP16 | $1.96 \cdot 10^1 \pm 0.27 \cdot 10^1$ | $2.28 \cdot 10^3 \pm 0.13 \cdot 10^3$ | $4.71 \cdot 10^3 \pm 0.52 \cdot 10^3$ | 6 |
| p65 | $2.23 \cdot 10^1 \pm 0.31 \cdot 10^1$ | $3.29 \cdot 10^3 \pm 0.20 \cdot 10^3$ | $8.50 \cdot 10^3 \pm 1.05 \cdot 10^3$ | 8.7 |
| VP64 | $1.55 \cdot 10^1 \pm 0.44 \cdot 10^1$ | $5.02 \cdot 10^3 \pm 0.44 \cdot 10^3$ | $7.33 \cdot 10^3 \pm 1.15 \cdot 10^3$ | 13.2 |
| VPR | $5.74 \pm 8.81$ | $2.25 \cdot 10^4 \pm 1.24 \cdot 10^4$ | $1.43 \cdot 10^4 \pm 0.93 \cdot 10^4$ | 59.2 |
| NFZ | $5.77 \pm 9.36$ | $1.38 \cdot 10^4 \pm 0.41 \cdot 10^4$ | $9.15 \cdot 10^3 \pm 3.64 \cdot 10^3$ | 36.3 |
| Binding domain: ZF43 | | | | |
| Activation domain | $\alpha_0$ (a.u.) | $\alpha$ (a.u.) | $K$ (a.u.) | $\beta_2$ (1 / s) |
| VP16 | $2.72 \cdot 10^1 \pm 0.24 \cdot 10^1$ | $5.42 \cdot 10^2 \pm 0.24 \cdot 10^2$ | $2.28 \cdot 10^3 \pm 0.28 \cdot 10^3$ | 1.4 |
| p65 | $2.49 \cdot 10^1 \pm 0.38 \cdot 10^1$ | $9.13 \cdot 10^2 \pm 0.92 \cdot 10^2$ | $3.56 \cdot 10^3 \pm 0.80 \cdot 10^3$ | 2.4 |
| VP64 | $2.63 \cdot 10^1 \pm 0.21 \cdot 10^1$ | $1.19 \cdot 10^3 \pm 0.08 \cdot 10^3$ | $5.79 \cdot 10^3 \pm 0.77 \cdot 10^3$ | 3.1 |
| VPR | $1.01 \cdot 10^1 \pm 0.77 \cdot 10^1$ | $1.20 \cdot 10^4 \pm 0.36 \cdot 10^4$ | $8.37 \cdot 10^3 \pm 3.16 \cdot 10^3$ | 31.6 |
| NFZ | $6.8 \pm 9.54$ | $1.96 \cdot 10^4 \pm 0.71 \cdot 10^4$ | $1.23 \cdot 10^4 \pm 0.56 \cdot 10^4$ | 51.7 |

**Table S3. List of plasmids used in transfection experiments.** Sequences in parenthesis are encoded in synthetic introns. Sequences in brackets are encoded in the antisense direction.

| Name | Plasmid ID | Description | Figures |
|---|---|---|---|
| pDRG007 | pKG4205 | EFS-mRuby2-ZF43-bGH | Figures S2 and S6B |
| pDRG008 | pKG4206 | EFS-mRuby2-ZF37-bGH | Figures S2 and S6B |
| pDRG011 | pKG4209 | ZF43x6-C-tagBFP-bGH | Figures S2 and S6B |
| pDRG012 | pKG4210 | ZF37x6-C-tagBFP-bGH | Figures S2 and S6B |
| pEP1.3.022 | pKG2568 | ZF37x6-C-VP16-ZF37-P2A-mGL-ZF43-bGH | Figures 2, 3, S3, 4, S4 and 6 |
| pEP1.3.023 | pKG2569 | ZF43x6-C-VP16-ZF43-P2A-mRuby2-ZF37-bGH | Figures 2, 3, S3, S4, 4 and 6 |
| pEP118 | pKG4158 | ZF37x6-C-mRuby2-bGH | Figures 4A, S6 and S7 |
| pEP119 | pKG4159 | EF1a-VP16-ZF43-P2A-mGL-bGH | Figures S2, 4A, S6 and S7 |
| pEP120 | pKG4160 | EF1a-VP16-ZF37-P2A-mGL-bGH | Figures S2, 4A, S6 and S7 |
| pEP121 | pKG4161 | EF1a-VP64-ZF43-P2A-mGL-bGH | Figures 4A, S6 and S7 |
| pEP122 | pKG4162 | EF1a-VP64-ZF37-P2A-mGL-bGH | Figures 4A, S6 and S7 |
| pEP123 | pKG4163 | EF1a-VPR-ZF43-P2A-mGL-bGH | Figures 4A, S6 and S7 |
| pEP124 | pKG4164 | EF1a-VPR-ZF37-P2A-mGL-bGH | Figures 4A, S6 and S7 |
| pEP125 | pKG4165 | EF1a-p65-ZF43-P2A-mGL-bGH | Figures 4A, S6 and S7 |
| pEP126 | pKG4166 | EF1a-p65-ZF37-P2A-mGL-bGH | Figures 4A, S6 and S7 |
| pEP127 | pKG4167 | EF1a-mGL(miR-FF6)-bGH | Figure S14 |
| pEP129 | pKG4169 | EF1a-mRuby2-TS.FF5x1-bGH | Figure S14 |
| pEP134 | pKG4174 | EF1a-mRuby2-TS.FF6x1-bGH | Figure S14 |
| pEP139 | pKG4186 | EF1a-mGL(miR-FF5)-bGH | Figure S14 |
| pEP150 | pKG4285 | ZF37x6-C-p65-ZF37-P2A-mGL-ZF43-bGH | Figures 4, 6 and S8 |
| pEP151 | pKG4286 | ZF43x6-C-p65-ZF43-P2A-mRuby2-ZF37-bGH | Figures 4, 6 and S8 |
| pEP154 | pKG4289 | ZF37x6-C-VPR-ZF37-P2A-mGL-ZF43-bGH | Figures 4, 5, 6, S10 and S12 |
| pEP155 | pKG4290 | ZF43x6-C-VPR-ZF43-P2A-mRuby2-ZF37-bGH | Figures 4, 5, 6, S10 and S13 |
| pEP202 | pKG4516 | ZF37x6-C-VP16-ZF37-P2A-mGL(miR-FF5)-ZF43-TS.FF6x1-bGH | Figures 6 and S15 |
| pEP203 | pKG4517 | ZF43x6-C-VP16-ZF43-P2A-mRuby2(miR-FF6)-ZF37-TS.FF5x1-bGH | Figures 6 and S15 |
| pEP204 | pKG4518 | EF1a-VP16-ZF37-P2A-mGL(miR-FF5)-ZF43-bGH | Figure 2 |
| pEP205 | pKG4519 | EF1a-VP16-ZF43-P2A-mRuby2(miR-FF6)-ZF37-bGH | Figure 2 |
| pEP206 | pKG4520 | EF1a-VP16-ZF37-P2A-mGL-ZF43-bGH | Figure S5 |
| pEP207 | pKG4521 | EF1a-VP16-ZF43-P2A-mRuby2-ZF37-bGH | Figure S5 |
| pEP232 | pKG4568 | ZF37x6-C-VP64-ZF37-P2A-mGL-ZF43-bGH | Figures 4, 6 and S9 |
| pEP233 | pKG4569 | ZF43x6-C-VP64-ZF43-P2A-mRuby2-ZF37-bGH | Figures 4, 6 and S9 |
| pEP234 | pKG4570 | ZF37x6-C-NFZ-ZF37-P2A-mGL-ZF43-bGH | Figures 4, 5, 6, S11 and S13 |

Continued on next page

Table S3 – continued from previous page

| Name | Plasmid ID | Description | Figures |
|---|---|---|---|
| pEP235 | pKG4571 | ZF43x6-C-NFZ-ZF43-P2A-mRuby2-ZF37-bGH | Figures 4, 5, 6, S11 and S12 |
| pEP236 | pKG4583 | ZF37x6-C-VP64-ZF37-P2A-mGL(miR-FF5)-ZF43-TS.FF6x1-bGH | Figures 6 and S17 |
| pEP237 | pKG4572 | ZF43x6-C-VP64-ZF43-P2A-mRuby2(miR-FF6)-ZF37-TS.FF5x1-bGH | Figures 6 and S17 |
| pEP238 | pKG4573 | ZF37x6-C-NFZ-ZF37-P2A-mGL(miR-FF5)-ZF43-TS.FF6x1-bGH | Figures 6 and S19 |
| pEP239 | pKG4574 | ZF43x6-C-NFZ-ZF43-P2A-mRuby2(miR-FF6)-ZF37-TS.FF5x1-bGH | Figures 6 and S19 |
| pEP240 | pKG4575 | ZF37x6-C-p65-ZF37-P2A-mGL(miR-FF5)-ZF43-TS.FF6x1-bGH | Figures 6 and S16 |
| pEP241 | pKG4584 | ZF43x6-C-p65-ZF43-P2A-mRuby2(miR-FF6)-ZF37-TS.FF5x1-bGH | Figures 6 and S16 |
| pEP242 | pKG4585 | ZF37x6-C-VPR-ZF37-P2A-mGL(miR-FF5)-ZF43-TS.FF6x1-bGH | Figures 6 and S18 |
| pEP243 | pKG4576 | ZF43x6-C-VPR-ZF43-P2A-mRuby2(miR-FF6)-ZF37-TS.FF5x1-bGH | Figures 6 and S18 |
| pEP405 | pKG4896 | ZF37x6-C-VP16-ZF37-P2A-mGL-ZF43-bGH-[ZF43x6-C-VP16-ZF43-P2A-mRuby2-ZF37-bGH] | Figure 7, convergent |
| pEP406 | pKG4897 | [ZF37x6-C-VP16-ZF37-P2A-mGL-ZF43-bGH]-ZF43x6-C-VP16-ZF43-P2A-mRuby2-ZF37-bGH | Figure 7, divergent |
| pEP407 | pKG4898 | ZF37x6-C-VP16-ZF37-P2A-mGL-ZF43-bGH-ZF43x6-C-VP16-ZF43-P2A-mRuby2-ZF37-bGH | Figure 7, tandem 1 |
| pEP408 | pKG4899 | ZF43x6-C-VP16-ZF43-P2A-mRuby2-ZF37-bGH-ZF37x6-C-VP16-ZF37-P2A-mGL-ZF43-bGH | Figure 7, tandem 2 |
| pEP409 | pKG4900 | ZF37x6-C-VP16-ZF37-P2A-mGL(miR-FF5)-ZF43-TS.FF6x1-bGH-[ZF43x6-C-VP16-ZF43-P2A-mRuby2(miR-FF6)-ZF37-TS.FF5x1-bGH] | Figure 7, convergent |
| pEP410 | pKG4901 | [ZF37x6-C-VP16-ZF37-P2A-mGL(miR-FF5)-ZF43-TS.FF6x1-bGH]-ZF43x6-C-VP16-ZF43-P2A-mRuby2(miR-FF6)-ZF37-TS.FF5x1-bGH | Figure 7, divergent |
| pEP411 | pKG4902 | ZF37x6-C-VP16-ZF37-P2A-mGL(miR-FF5)-ZF43-TS.FF6x1-bGH-ZF43x6-C-VP16-ZF43-P2A-mRuby2(miR-FF6)-ZF37-TS.FF5x1-bGH | Figure 7, tandem 1 |
| pEP412 | pKG4903 | ZF43x6-C-VP16-ZF43-P2A-mRuby2(miR-FF6)-ZF37-TS.FF5x1-bGH-ZF37x6-C-VP16-ZF37-P2A-mGL(miR-FF5)-ZF43-TS.FF6x1-bGH | Figure 7, tandem 2 |
| pEP423 | pKG5333 | ZF37x6-C-NFZ-ZF37-P2A-mGL-ZF43-bGH-[ZF43x6-C-NFZ-ZF43-P2A-mRuby2-ZF37-bGH] | Figure 7, convergent |
| pEP424 | pKG5334 | [ZF37x6-C-NFZ-ZF37-P2A-mGL-ZF43-bGH]-ZF43x6-C-NFZ-ZF43-P2A-mRuby2-ZF37-bGH | Figure 7, divergent |
| pEP425 | pKG5335 | ZF37x6-C-NFZ-ZF37-P2A-mGL-ZF43-bGH-ZF43x6-C-NFZ-ZF43-P2A-mRuby2-ZF37-bGH | Figure 7, tandem 1 |

Continued on next page

Table S3 – continued from previous page

| Name | Plasmid ID | Description | Figures |
|---|---|---|---|
| pEP426 | pKG5336 | ZF43x6-C-NFZ-ZF43-P2A-mRuby2-ZF37-bGH-ZF37x6-C-NFZ-ZF37-P2A-mGL-ZF43-bGH | Figure 7, tandem 2 |
| pEP427 | pKG5337 | ZF37x6-C-NFZ-ZF37-P2A-mGL(miR-FF5)-ZF43-TS.FF6x1-bGH-[ZF43x6-C-NFZ-ZF43-P2A-mRuby2(miR-FF6)-ZF37-TS.FF5x1-bGH] | Figure 7, convergent |
| pEP428 | pKG5338 | [ZF37x6-C-NFZ-ZF37-P2A-mGL(miR-FF5)-ZF43-TS.FF6x1-bGH]-ZF43x6-C-NFZ-ZF43-P2A-mRuby2(miR-FF6)-ZF37-TS.FF5x1-bGH | Figure 7, divergent |
| pEP429 | pKG5339 | ZF37x6-C-NFZ-ZF37-P2A-mGL(miR-FF5)-ZF43-TS.FF6x1-bGH-ZF43x6-C-NFZ-ZF43-P2A-mRuby2(miR-FF6)-ZF37-TS.FF5x1-bGH | Figure 7, tandem 1 |
| pEP430 | pKG5340 | ZF43x6-C-NFZ-ZF43-P2A-mRuby2(miR-FF6)-ZF37-TS.FF5x1-bGH-ZF37x6-C-NFZ-ZF37-P2A-mGL(miR-FF5)-ZF43-TS.FF6x1-bGH | Figure 7, tandem 2 |
| pGEEC510 | pKG2116 | EF1a-mRuby2-bGH | Figure S14 |
| pGEEC525 | pKG2215 | ZF43x6-C-mRuby2-bGH | Figures 4A, S6 and S7 |
| pGEEC549 | pKG2898 | EF1a-tagBFP-bGH | Figures 3, S3, S4, 4, S5, 5, S6, 6, S7, S8, S9, S10, S11, S12, S13, S15, S16, S17, S18 and S19 |
| pGEEC555 | pKG2908 | EF1a-SNAP-iRFP.d-bGH | Figures 2, S2, 3, S3, S4, 4, S5, 5, 6, S6B, 7, S8, S9, S10, S11, S12, S13, S15, S16, S17, S18 and S19 |
| pMJ001 | pKG4539 | EF1a-NFZ-ZF43-P2A-mGL-bGH | Figures 4A, S6 and S7 |
| pMJ002 | pKG4540 | EF1a-NFZ-ZF37-P2A-mGL-bGH | Figures 4A, S6 and S7 |

**Table S4. Attune NxT flow cytometer channels and voltages**

| Channel | Laser (nm) | Filter | PMT Voltage |
|---|---|---|---|
| Figures S2 and S6B | | | |
| tagBFP | 405 | 440 / 50 | 220 |
| mGL | 488 | 510 / 10 | 220 |
| mRuby2 | 561 | 615 / 25 | 300 |
| iRFP670 | 637 | 670 / 14 | 280 |
| FSC | | | 100 |
| SSC | | | 360 |
| All figures unless otherwise noted | | | |
| tagBFP | 405 | 440 / 50 | 220 |
| mGL | 488 | 530 / 30 | 220 |
| mRuby2 | 561 | 620 / 15 | 300 |
| iRFP670 | 637 | 670 / 14 | 280 |
| FSC | | | 100 |
| SSC | | | 360 |
